# Imprinted gene expression at the *Dlk1-Dio3* cluster is controlled by both maternal and paternal *IG-DMR*s in a tissue-specific fashion

**DOI:** 10.1101/536102

**Authors:** Katherine A. Alexander, María J. García-García

## Abstract

Imprinting at the *Dlk1-Dio3* cluster is controlled by the *IG-DMR*, an imprinting control region differentially methylated between maternal and paternal chromosomes. The maternal *IG-DMR* is essential for imprinting control, functioning as a *cis* enhancer element. Meanwhile, DNA methylation at the paternal *IG-DMR* is thought to prevent enhancer activity. To explore whether suppression of enhancer activity at the methylated *IG-DMR* requires the transcriptional repressor TRIM28, we analyzed *Trim28^chatwo^* embryos and performed epistatic experiments with *IG-DMR* deletion mutants. We found that while TRIM28 regulates the enhancer properties of the paternal *IG-DMR*, it also controls imprinting through other mechanisms. Additionally, we found that the paternal *IG-DMR*, previously deemed dispensable for imprinting, is required in certain tissues, demonstrating that imprinting is regulated in a tissue-specific manner. Using PRO-seq to analyze nascent transcription, we identified 30 novel transcribed regulatory elements, including 23 that are tissue-specific. These results demonstrate that different tissues have a distinctive regulatory landscape at the *Dlk1-Dio3* cluster and provide insight into potential mechanisms of tissue-specific imprinting control. Together, our findings challenge the premise that *Dlk1-Dio3* imprinting is regulated through a single mechanism and demonstrate that different tissues use distinct strategies to accomplish imprinted gene expression.

## INTRODUCTION

Genomic imprinting is an epigenetic mechanism that controls the allele-specific expression of certain genes depending on their maternal or paternal inheritance. Mutations disrupting imprinted gene expression result in embryonic lethality in mice, congenital defects in humans, and affect the reprogramming of induced pluripotent stem cells, highlighting the importance of imprinting control for embryonic development and cell differentiation (1, 2). Most imprinted genes reside in gene clusters, where their expression is controlled by imprinting control regions (ICRs), regulatory sequences that are differentially methylated between the maternally and paternally inherited chromosomes (3). DNA methylation at these differentially methylated regions (DMRs) has been proposed to influence the allele-specific recruitment of transcription factors that function in *cis* to control gene expression. However, while a few proteins are known to bind certain ICRs on a methylation-dependent fashion (4-15), the information to date suggests that there is not a universal mechanism of imprinting control. Instead, epigenetic marks at ICRs are interpreted in a manner that is specific for each imprinted cluster (reviewed in 3).

While most imprinted genes are expressed in an allele-specific fashion in all tissues, others show more dynamic imprinting control mechanisms. For instance, as many as 28% of all known imprinted genes display tissue-specific imprinting, with monoallelic expression in only one or a few tissues, and biallelic expression in all others (16-18). Other genes are imprinted in a stage-specific manner, being biallelically expressed early in development and only becoming monoallelically expressed at later embryonic stages, or vice versa (19-21). Finally, a few imprinted genes undergo imprinting reversal in a tissue-specific or stage-specific fashion, with expression from the maternal allele in some tissues or developmental stages, and from the paternal allele in others (22-24). Despite their dynamic allele-specific expression, many genes subject to tissue and/or stage-specific imprinting still preserve the characteristic allele-specific DNA methylation at their ICRs (reviewed in 17). Therefore, either epigenetic marks at ICRs are interpreted in a context-specific fashion, or other regulatory sequences at imprinted clusters provide additional layers of tissue and stage-specific imprinting control (23, 25-32).

The *Dlk1-Dio3* imprinted gene cluster contains three protein-coding genes, *Dlk1*, *Rtl1* and *Dio3*, which are expressed from the paternal allele, and several non-coding RNAs, including *Gtl2*, *Rtl1as*, *Rian* and *Mirg*, that are expressed from the maternal allele as a single transcriptional unit (Figure 1A; reviewed in 33). The allele-specific expression of these genes is maintained in most tissues analyzed (18). Data from mouse embryos and different human tissues has failed to provide evidence for tissue-specific imprinting of *Gtl2* (18, 34). However, *Dlk1* is biallelically expressed in astrocytes (35), shows a “relaxation” of imprinting in the liver (15% expression from the maternal allele; 36) and is subject to imprinting reversal in ESCs (24), while *Dio3* is biallelically expressed in the placenta and a few other tissues (37). Additionally, imprinted genes in the *Dlk1-Dio3* are widely expressed during embryogenesis, but the levels of expression of *Gtl2* and *Dlk1* are variable across tissues and developmental stages (35, 38-42).

**Figure 1.**
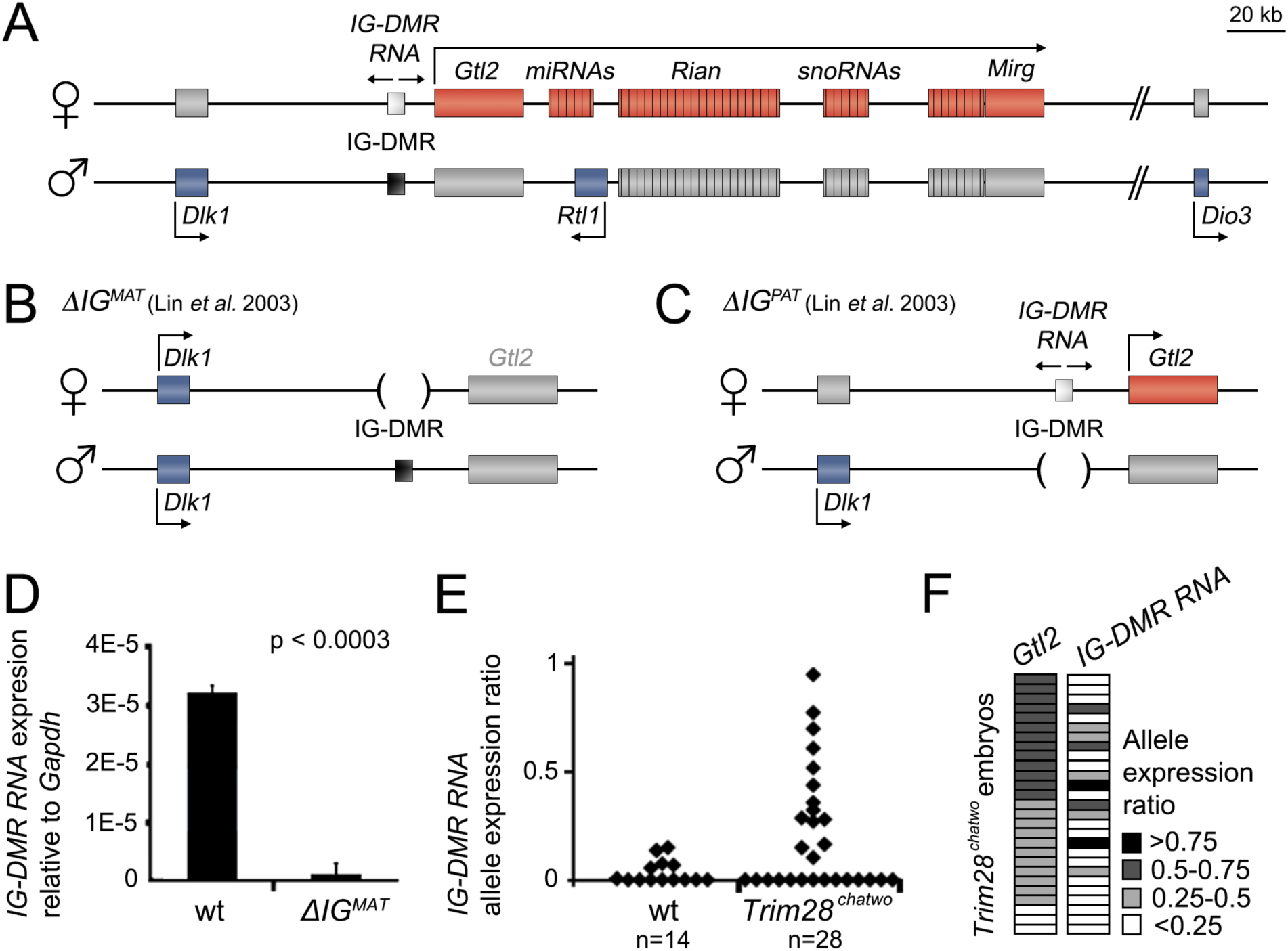
*IG-DMR RNA* expression in *Trim28^chatwo^* embryos. **(A)** Schematic representation of the *Dlk1-Dio3* locus. Colored boxes denote genes with maternal (red), paternal (blue) or no expression (grey). *The IG-DMR* (small rectangle) is methylated (black) on the paternal allele. Arrows indicate the direction of transcription at gene promoters, as well as bidirectional transcription at the *IG-DMR*. **(B-C)** Illustration of the effects of the maternal (*ΔIG^MAT^*) and paternal (*ΔIG^PAT^*) *IG-DMR* deletion, as reported in (31). **(D)** qRT-PCR analysis of *IG-DMR RNA* expression relative to *Gapdh* in wild type and *ΔIG^MAT^* E8.5 embryos. Bars show the average of three individual embryos. Two technical replicates were analyzed for each biological sample. Error bars represent standard deviation. p-value was calculated using Student’s unpaired T-test. **(E)** Allele expression ratio of *IG-DMR RNA* in individual wild type and *Trim28^chatwo^* embryos measured by Sanger sequencing and quantified with PeakPicker. Allele ratios were calculated as paternal/maternal expression (0 = monoallelic maternal expression, 1 = equal expression from both parental alleles). **(F)** Plot indicating the allele expression ratios of *Gtl2* and *IG-DMR RNA* in individual *Trim28^chatwo^* embryos. Each line represents a single embryo. Embryos are sorted by decreasing *Gtl2* allele expression ratio. *IG-DMR RNA* primer location is shown in Supplementary Figure 1.

Imprinted gene expression within the *Dlk1-Dio3* cluster is controlled by an InterGenic Differentially Methylated Region (*IG-DMR*) located between *Dlk1* and *Gtl2* (43). The maternal *IG-DMR* is unmethylated and has been shown to be required for *cis* activation of maternal genes, as well as for *cis* repression of paternal genes in embryonic tissues (31). A 4.15 kb deletion of the maternal *IG-DMR* allele in mice is embryonic lethal and causes loss of *Gtl2* expression, as well as biallelic expression of *Dlk1* (Figure 1B; 31). In contrast, the same 4.15 kb deletion of the methylated paternal *IG-DMR* allele does not disrupt imprinting and produces adult mice without discernible phenotypes, suggesting that the paternal *IG-DMR* is dispensable for imprinting control (Figure 1C; 31, 44). Loss of methylation at the paternal *IG-DMR* causes a maternalization of the paternal chromosome, with biallelic expression of *Gtl2* and biallelic repression of *Dlk1* (38, 45). Therefore, the evidence to date indicates that the unmethylated maternal *IG-DMR* is functionally important to regulate imprinted gene expression in *cis*, while DNA methylation prevents transcriptional regulatory activity at the paternal *IG-DMR*. This model is further supported by recent studies identifying maternal-specific enhancer-like properties within the *IG-DMR*, including transcription of bidirectional short non-coding “enhancer RNAs” (eRNAs) and enhancer-specific histone modifications (24, 46, 47), as well as interactions between the *IG-DMR* and the *Gtl2* promoter (46). Supporting the notion that DNA methylation abrogates the enhancer activity of the *IG-DMR*, the paternal methylated *IG-DMR* did not show enhancer-like properties in mESCs (24). Additionally, a recent study using a small 420bp *IG-DMR* deletion identified a CpG tandem repeat region within the *IG-DMR* that is required paternally to maintain *IG-DMR* methylation, but is dispensable for imprinting control when inherited maternally, suggesting that the dual roles of the *IG-DMR* to maintain paternal imprints and to function as a cis-enhancer element are physically separable, yet functionally connected (48).

DNA methylation at the paternal *IG-DMR* is known to influence the allele-specific recruitment of certain transcriptional regulators such as ZFP57 and TRIM28, which form a transcriptional repressive protein complex (4, 45, 49). Deletion of either *Zfp57* or *Trim28* during early development results in biallelic *Gtl2* expression, supporting an important role for these transcription factors in regulating *Dlk1-Dio3* imprinting (45, 50, 51). Because loss of *Gtl2* imprinting in maternal-zygotic *Zfp57* mutants and *Trim28* null embryos is accompanied by loss of DNA methylation at the paternal *IG-DMR*, it has been proposed that ZFP57-TRIM28 complexes regulate imprinting by maintaining methyl marks at germline DMRs during the genome-wide reprogramming events that take place shortly after fertilization (45, 50). However, the analysis of conditional *Trim28* mutants, as well as hypomorphic *Trim28^chatwo^* embryos, showed that loss of *Trim28* at later embryonic stages disrupts *Gtl2* imprinting without affecting the methylation status of the *IG-DMR*, providing evidence that TRIM28 controls imprinting after genome-wide reprograming through molecular mechanisms that are distinct from its early role to preserve DNA methylation (50).

To gain mechanistic insight into how gene expression is regulated at the *Dlk1-Dio3* cluster, we first inquired whether TRIM28 functions after genome-wide reprogramming by binding to the methylated *IG-DMR* and preventing paternal *IG-DMR* enhancer activity. By analyzing *IG-DMR RNA* expression in hypomorphic *Trim28^chatwo^* embryos and performing epistasis experiments with a deletion of the *IG-DMR*, we determined that TRIM28 regulates *Dlk1-Dio3* imprinting after genome-wide reprogramming through mechanisms that are unrelated to its ability to bind the methylated *IG-DMR*. During the course of these studies, we unexpectedly found that the paternal *IG-DMR*, previous deemed unnecessary for imprinting control, was required to repress paternal *Gtl2* expression in some embryonic tissues. Further analysis of *IG-DMR* deletion mutants revealed that both the maternal and paternal *IG-DMR*s are required for imprinting control, and that they perform different regulatory functions in a tissue-specific manner. These findings, together with results from a survey of regulatory sequences using PRO-seq, demonstrate that the control of imprinted gene expression at the *Dlk1-Dio3* cluster is strikingly dynamic, comprising multiple regulatory sequences that, together with germline imprints, function in a tissue-specific manner to accomplish imprinted gene expression.

## MATERIALS AND METHODS

### Mice

Experiments involving *IG-DMR RNA* allele-specific analysis in *Trim28^chatwo^* mutants used crosses between *Trim28^chatwo^* mice in a congenic FvB/NJ or C57BL/6J genetic background and *Trim28^chatwo^* mice raised in a mixed background that was predominantly FvB/NJ, but selected to be homozygous for CAST/EiJ at the *Dlk1-Dio3* cluster on chromosome 12 (FvB/CAST-12). *Trim28^chatwo^* embryos derived from these crosses arrest at E8.5 (52). The *Δ*IG-DMR deletion was generated in a 129/OlaHsd background (31) and maintained in a congenic C57BL/6J background. For *Δ*IG-DMR; *Trim28^chatwo^* epistasis experiments, *Δ*IG-DMR;*Trim28^chatwo^* males were crossed with *Trim28^chatwo^* mice from the mixed FvB/CAST-12 background, which generated littermate embryos of each genotype analyzed at E8.5 in Figure 2. For experiments involving allele-specific analysis in E14.5 *Δ*IG-DMR mutants, *Δ*IG-DMR deletion males (for *Δ*IG^PAT^) or females (*Δ*IG^MAT^) were mated to wild type mice from the mixed FvB/CAST-12 genetic background. Genotyping primers for *ΔIG-DMR*, *Trim28^chatwo^* and for polymorphisms at chromosome 12 are shown in Supplementary Table 1.

**Figure 2.**
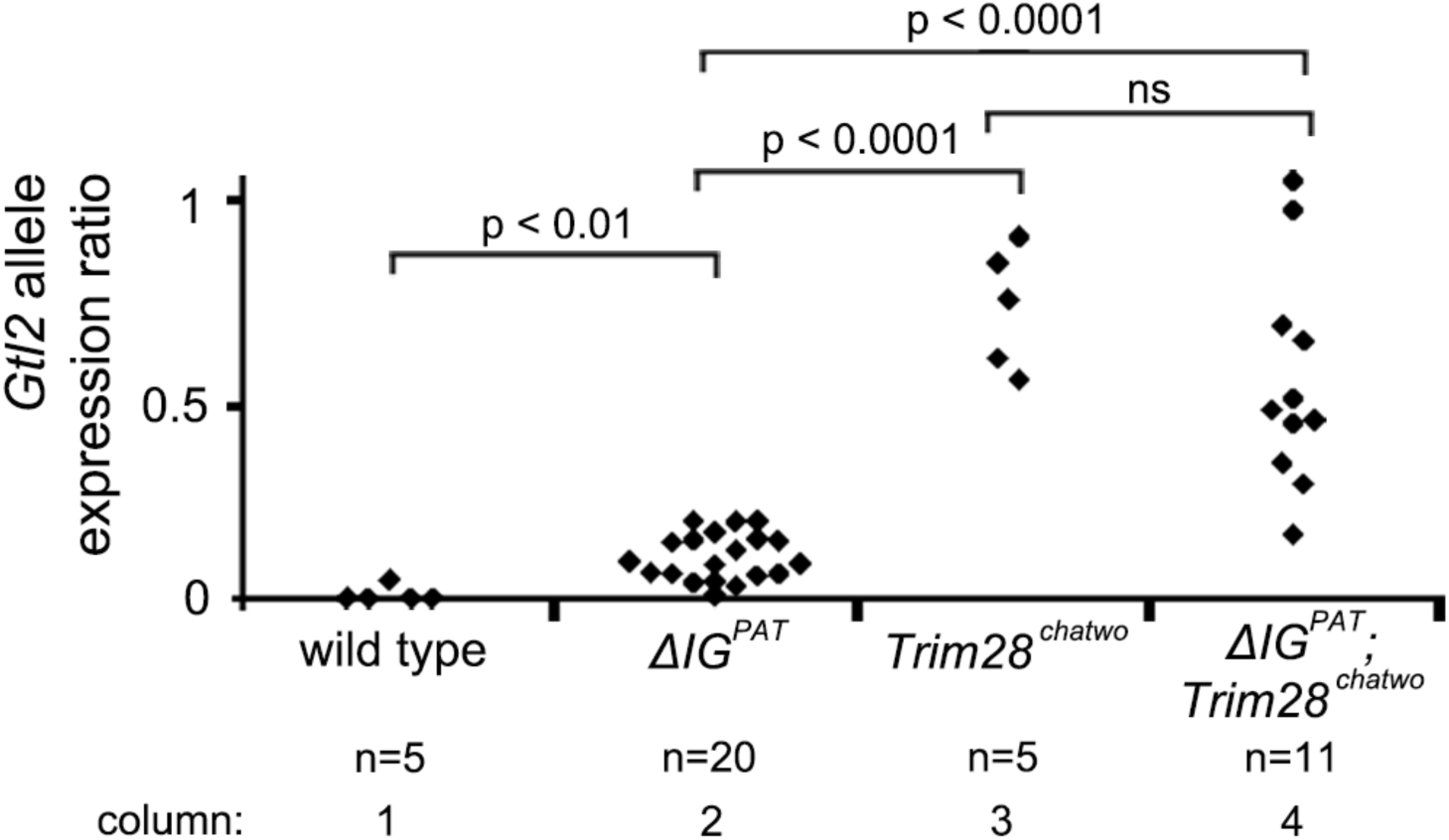
*Gtl2* allele expression ratio in wild type, *ΔIG^PAT^, Trim28^chatwo^* and *ΔIG^PAT^;Trim28^chatwo^* E8.5 embryos. Allele expression was measured in individual embryos bearing allele-specific polymorphisms using Sanger sequencing and was quantified as a ratio of paternal/maternal expression by PeakPicker (0 = monoallelic maternal expression, 1 = equal expression from both parental alleles). p-values were calculated using Student’s unpaired T-test. ns-not significant.

### Embryo collection

Embryos were dissected in ice cold phosphate buffered saline with 4% bovine serum albumin. For analysis of individual E14.5 tissues, the yolk sac, liver, lung and limb buds were dissected from the embryo and flash-frozen in liquid nitrogen. For E8.5 embryos, the yolk sac was dissected from embryonic tissues and analyzed separately.

### Expression analysis

RNA was isolated from mouse embryonic tissues using RNA-STAT60. RNA was DNase I-treated prior to First Strand cDNA synthesis using SuperScript III with random hexamer priming (Invitrogen). Expression was analyzed by quantitative reverse-transcriptase amplification (qRT-PCR) using KleenGreen (IBI). Expression was normalized to *Gapdh* or *β*-actin as indicated. The median amongst technical replicates was calculated to quantify each biological sample.

Allele specific expression was assayed by Sanger sequencing of SNP-containing RT-PCR products and quantified as the ratio of paternal:maternal expression using PeakPicker (53). For primer sequences see Supplementary Table 1. Statistical significance was evaluated by unpaired two-tailed Student’s T-tests.

### PRO-seq and dREG

Liver and yolk sac from six E14.5 embryos were flash-frozen, then broken up into a fine powder in liquid nitrogen with a pestle. Chromatin from these extracts was isolated and stored as described (http://biorxiv.org/lookup/doi/10.1101/185991). Precision Nuclear Run-On was performed on chromatin samples and then processed for sequencing as described (54). PRO-seq was performed on two replicates for each tissue. Samples were obtained from embryos containing parent-of-origin polymorphisms at the *Dlk1-Gtl2* locus on Chr. 12 (C57BL/6J, maternal; CAST/EiJ, paternal). dREG was performed as previously described (55) on each PRO-seq dataset. Only common dREG peaks between replicates were considered. A validation of dREG results was performed by comparing identified peaks with ENCODE ChIP data on histone marks characteristic of enhancers (H3K27ac and H3K4me1) and promoters (H3K27ac and H3K4me3). See Supplementary Figure 6 for details. PRO-seq data was deposited in the NCBI GEO database under the accession number GSE119882.

**Figure 6.**
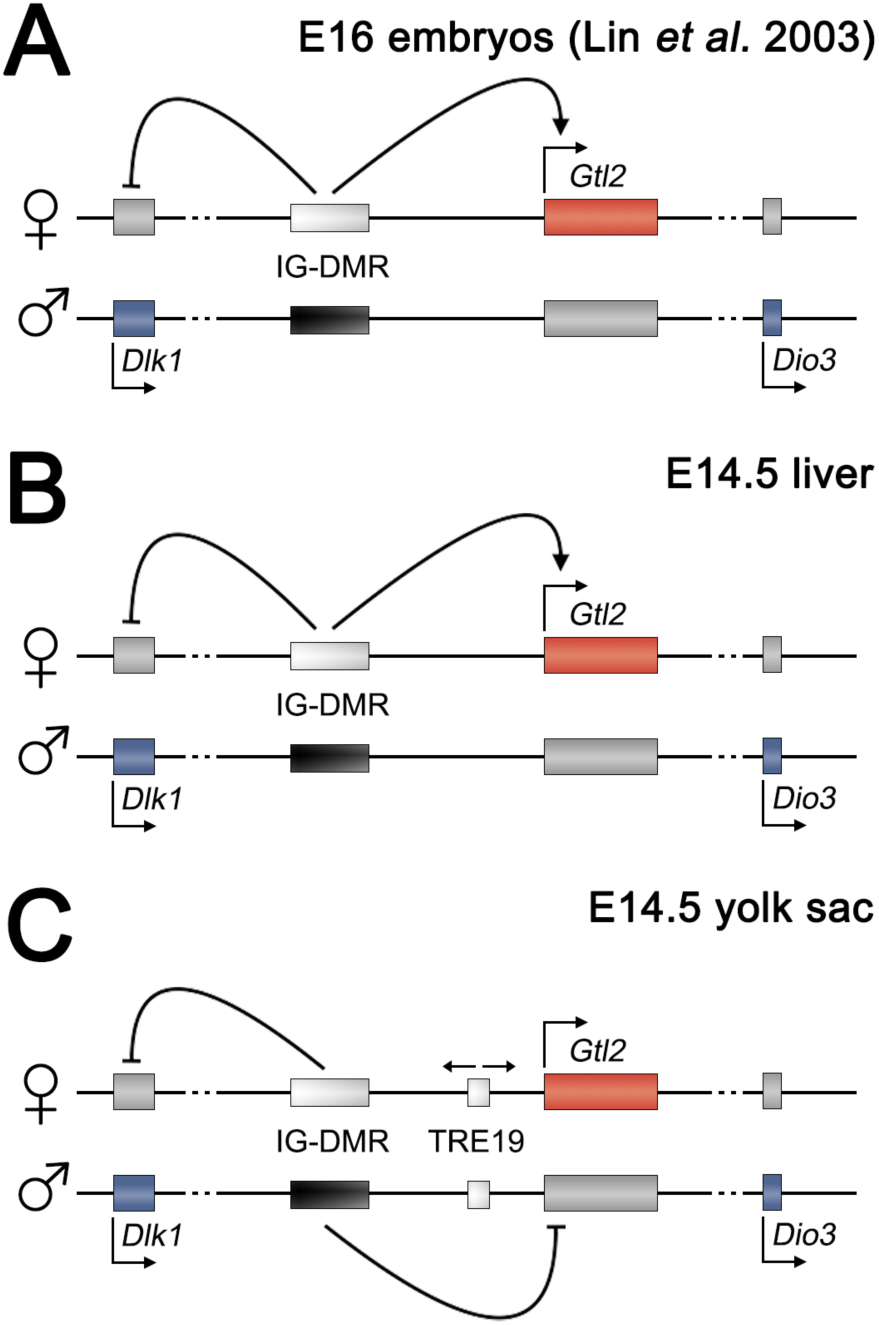
Tissue-specific imprinting control at the *Dlk1-Dio3* cluster. Schematic views of the *Dlk1-Dio3* cluster indicating the regulatory relationships of the maternal and the paternal *IG-DMRs* in E16 embryos (31) **(A)**, E14.5 liver **(B)**, and E14.5 yolk sac **(C)**. Pointed arrows indicate a positive regulatory relationship and blunted arrows a repressive role. Colored boxes denote genes with maternal (red), paternal (blue) or no expression (grey). Small arrows indicate the direction of transcription at gene promoters, as well as bidirectional transcription at TRE19. The locations of transcripts and regulatory sequences is not drawn to scale.

## RESULTS

### The paternal *IG-DMR* displays enhancer-like properties in *Trim28* hypomorphic mutants

Our previous analysis showed that TRIM28 controls genomic imprinting through distinct mechanisms during and after genome-wide reprogramming. Specifically, we found that conditional *Trim28* mutants and hypomorphic *Trim28^chatwo^* embryos had biallelic *Gtl2* expression without affecting *IG-DMR* methylation, providing evidence that TRIM28 functions through mechanisms that are distinct from its early role maintaining DNA methylation at germline imprints during genome-wide reprogramming (50). To investigate the mechanisms used by TRIM28 to control imprinting after genome-wide reprogramming, we first tested whether TRIM28 regulates the enhancer properties of the *IG-DMR*. To this end, we analyzed E8.5 embryos homozygous for the *Trim28^chatwo^* allele, a condition that decreases the transcriptional repressive activity and protein stability of TRIM28 (52), and used expression of RNA from the *IG-DMR* (hereby called *IG-DMR RNAs*) as a proxy for enhancer activity (24, 56-58). Because previous studies showing *IG-DMR RNA* expression were performed in mESCs (24), we first sought to confirm whether maternal-specific enhancer-like transcription could be observed in embryonic samples. To measure transcription from the *IG-DMR*, we performed qRT-PCR, using embryos with a maternal 4.15 kb *IG-DMR* deletion (*Δ*IG^MAT^; 31) as negative controls. We found that *IG-DMR RNA* was expressed in E8.5 wild type embryos at very low levels with respect to *Gapdh*, but that these levels were statistically different compared to negative control *ΔIG^MAT^* embryos (Figure 1D, p < 0.0003). This result provides evidence that, similar to mESCs, *IG-DMR RNAs* are transcribed from the maternal allele in E8.5 embryos. To further confirm that embryonic *IG-DMR RNA* expression is maternal-specific, we sequenced RT-PCR products from E8.5 embryos that contained single nucleotide polymorphisms (SNPs) distinguishing the maternal and paternal alleles. We found that the paternal-to-maternal *IG-DMR RNA* allele expression ratios in individual E8.5 wild type embryos were very low (Figure 1E), demonstrating that *IG-DMR RNAs* are primarily expressed from the maternal allele. In contrast, analysis of allele-specific expression in *Trim28^chatwo^* embryos showed a significant number of embryos with high levels of *IG-DMR RNA* biallelic expression (Figure 1E), supporting a role for TRIM28 in repressing *IG-DMR RNA* expression from the paternal allele. Because *Trim28^chatwo^* embryos have wild type levels of DNA methylation at the *IG-DMR* (50), our findings suggest that TRIM28 regulates genomic imprinting after genome-wide reprogramming by preventing *IG-DMR* enhancer activity from the methylated *IG-DMR*.

### TRIM28 controls *Gtl2* imprinted expression through mechanisms other than binding the paternal *IG-DMR*

Our analysis of *IG-DMR* transcription in *Trim28^chatwo^* mutants showed variable levels of biallelic *IG-DMR RNA* expression between individual E8.5 embryos, with 14/28 *Trim28^chatwo^* mutants showing maternal-specific *IG-DMR RNA* expression and 14/28 others showing expression from both the maternal and paternal allele (Figure 1E). This partially-penetrant biallelic expression from the *IG-DMR* was reminiscent of the partially penetrant loss of *Gtl2* imprinting that we previously observed in *Trim28^chatwo^* mutants (50). To determine if biallelic expression of *Gtl2* in *Trim28^chatwo^* embryos could be explained by abnormal paternal *IG-DMR* enhancer activity, we measured *IG-DMR RNA* and *Gtl2* biallelic expression in single *Trim28^chatwo^* mutants. We found that all the *Trim28^chatwo^* embryos that biallelically expressed *IG-DMR RNA* also showed *Gtl2* biallelic expression (Figure 1F). However, the degree of biallelic *Gtl2* expression in these *Trim28^chatwo^* mutants did not correlate with the levels of *IG-DMR RNA* transcription. Additionally, many *Trim28^chatwo^* embryos showed biallelic *Gtl2* expression without biallelically expressing *IG-DMR RNA.* Therefore, while aberrant enhancer-like transcription at the paternal *IG-DMR* could contribute to *Trim28^chatwo^* imprinting defects, this ectopic enhancer-like activity cannot fully explain loss of *Gtl2* imprinting in *Trim28^chatwo^* mutants.

Previous studies suggest that TRIM28 regulates imprinting by binding to the methylated paternal *IG-DMR* (4). However, given that neither loss of *IG-DMR* DNA methylation (50), nor paternal *IG-DMR* enhancer-like activity (Figure 1F) can fully explain loss of imprinting in *Trim28^chatwo^* mutants, we hypothesized that TRIM28 might regulate *Gtl2* through alternative mechanisms that do not require TRIM28 binding to the paternal *IG-DMR*. To test this hypothesis, we performed epistasis experiments between *Trim28^chatwo^* mutants and mutants with the 4.15 kb deletion of the paternal *IG-DMR* (*Δ*IG^PAT^; 31). If TRIM28 controls imprinting at *Gtl2* solely by binding the paternal *IG-DMR*, we would expect the paternal *IG-DMR* deletion to be epistatic over loss of *Trim28.* In this scenario, *ΔIG^PAT^;Trim28^chatwo^* mutants would show maternal-specific expression of *Gtl2,* as previously reported for *Δ*IG^*PAT*^ mutants (Figure 1C; 31). In contrast to this expectation, we observed that *ΔIG^PAT^;Trim28^chatwo^* embryos expressed *Gtl2* biallelically and at similar levels to those found in *Trim28^chatwo^* mutants (Figure 2, columns 3-4), revealing that the *Trim28^chatwo^* mutation is epistatic over deletion of the paternal *IG-DMR*. These findings demonstrate that, to maintain proper *Gtl2* imprinting, TRIM28 must function by binding sequences other than the paternal *IG-DMR*.

### The paternal *IG-DMR* has tissue-specific requirements for imprinted gene expression

Deletion of the paternal *IG-DMR* has been previously shown to be dispensable for imprinting control in whole E16 embryos (Figure 1C; 31). Intriguingly, however, our analysis of *Δ*IG^*PAT*^ E8.5 embryos showed that although these mutants expressed *Gtl2* primarily from the maternal allele, there was a slight expression of paternal *Gtl2* compared to wild type littermates (Figure 2, column 1-2, p < 0.01). Because of the low levels of paternal *Gtl2* expression in *Δ*IG^*PAT*^ E8.5 embryos, we hypothesized that the paternal *IG-DMR* could be required to repress *Gtl2* only in a subset of tissues. To investigate possible tissue-specific functions of the paternal *IG-DMR*, we used qRT-PCR to test expression of *Dlk1*, *Gtl2* and *Dio3* in *Δ*IG^*PAT*^ embryos at E14.5, a developmental stage at which individual tissues can be reliably separated by dissection.

Analysis of *Dlk1-Dio3* imprinting in the E14.5 liver did not show any significant effects for the paternal *IG-DMR* deletion on either the expression of maternally-expressed *Gtl2*, or paternally-expressed *Dlk1* and *Dio3* (Figure 3A). Additionally, analysis of allele-specific expression of *Gtl2* using Sanger sequencing revealed that in wild type liver samples *Gtl2* was exclusively expressed from the maternal allele and that there were no statistically significant differences in *Δ*IG^*PAT*^ mutants (Figure 3C, wt n=6, *Δ*IG^*PAT*^ n=7). Therefore, the paternal *IG-DMR* is not required to regulate *Gtl2* expression in *cis* and is dispensable for imprinting control in the liver.

**Figure 3.**
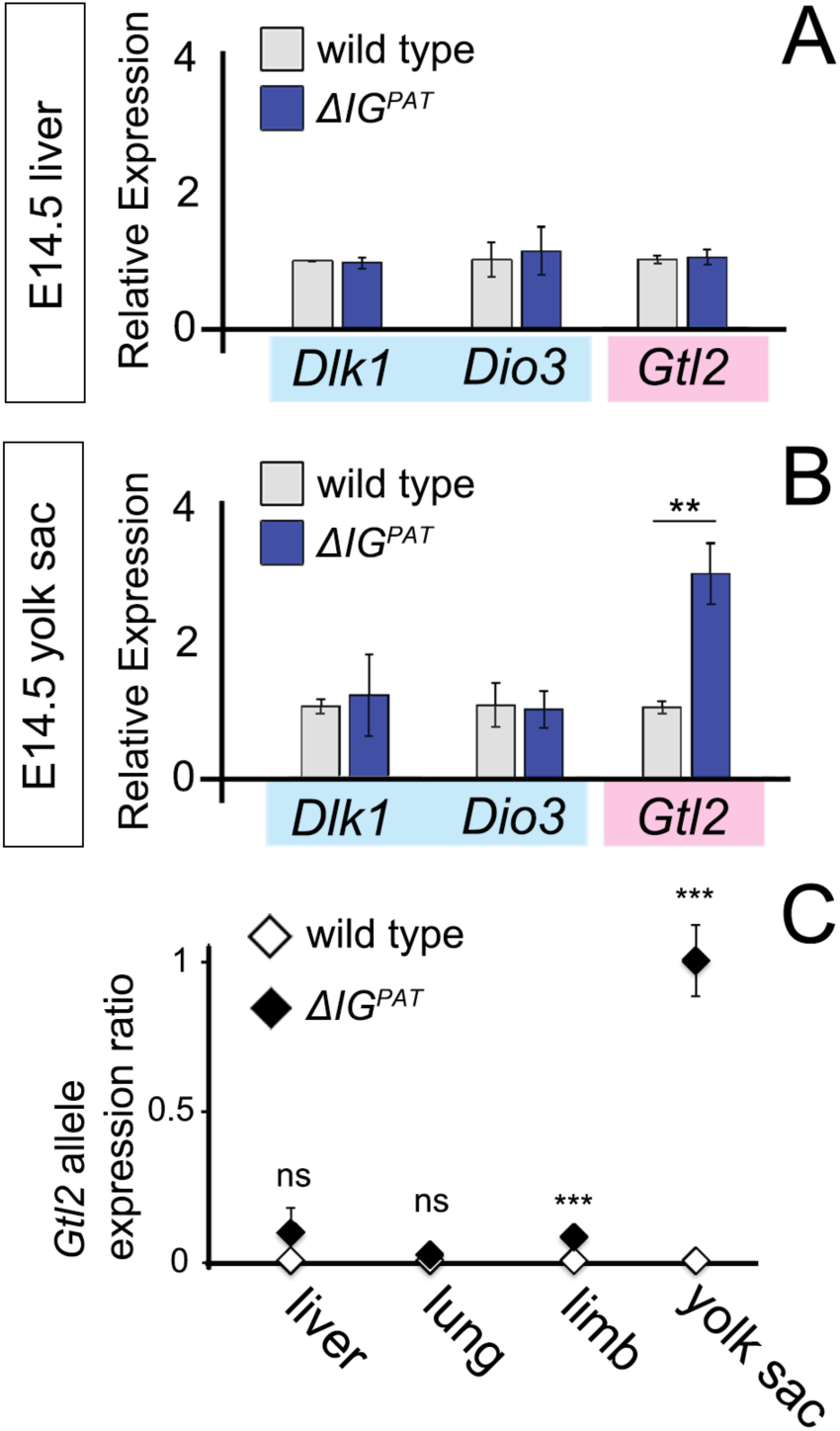
Tissue-specific effects of *ΔIG^PAT^* on *Dlk1-Dio3* gene expression. **(A-B)** qRT-PCR of paternally-expressed (blue, *Dlk1* and *Dio3*) and maternally-expressed (pink, *Gtl2*) genes in *ΔIG^PAT^* mutants (blue bars) in E14.5 liver **(A)**, and in E14.5 yolk sac **(B)** samples as compared to wild type controls (grey bars). Plot represents the average of three individual embryos. Expression is normalized to *β-actin* and relative to wild type littermates. Error bars represent SEM. **(C)** *Gtl2* allele expression ratio as measured by Sanger sequencing and quantified by PeakPicker in wild type and *ΔIG^PAT^* mutants. Results shown represent the average from six wild type and seven *ΔIG^PAT^* individual embryos. Three technical replicates were analyzed for each biological sample. ns-not significant, ** p < 0.01, *** p < 0.005

In contrast, we found that the paternal *IG-DMR* was required for imprinting control in the E14.5 yolk sac. In E14.5 yolk sacs from *Δ*IG^*PAT*^ mutants, *Dlk1* and *Dio3* expression levels were normal, but *Gtl2* expression was elevated compared to wild type littermate controls (Figure 3B), suggesting that the paternal *IG-DMR* is required for *cis* repression of *Gtl2* in the yolk sac. To further test a possible tissue-specific role of the paternal *IG-DMR* as a repressive *cis* regulatory element, we measured allelic expression of *Gtl2* in *Δ*IG^*PAT*^ samples. In contrast with liver, lung and limb tissues, where the *Δ*IG^*PAT*^ mutation did not have drastic effects on *Gtl2* allele-specific expression, we found that deletion of the paternal *IG-DMR* in E14.5 yolk sacs resulted in complete loss of *Gtl2* imprinting, with equal levels of *Gtl2* expression from the maternal and paternal alleles (Figure 3C, column 4, p < 0.005, wt n=6, *Δ*IG^*PAT*^ n=7). Therefore, our results show that the paternal *IG-DMR* is essential for silencing paternal *Gtl2* specifically in the yolk sac.

Overall, our analysis of *Δ*IG^*PAT*^ mutants demonstrates novel functions of the paternal *IG-DMR* and reveals that this regulatory sequence is required for imprinting control in a tissue-specific manner.

### Tissue-specific requirements for the maternal *IG-DMR*

A previous study showed that deletion of the maternal *IG-DMR* has different effects on *Dlk1-Dio3* imprinted gene expression in E16 placentas, as compared with their respective embryonic samples. Specifically, the maternal *IG-DMR*, which is required in embryos for expression of both maternally and paternally expressed genes in the *Dlk1-Dio3* cluster (Figure 1B; 31), was found to be dispensable for *Gtl2* expression in *ΔIG^MAT^* placentas (44). These findings prompted us to evaluate whether the maternal *IG-DMR* also has specific requirements in other tissues. To this end, we analyzed *Dlk1-Dio3* gene expression by qRT-PCR in the yolk sac and liver of *ΔIG^MAT^* mutants.

We found that the maternal *IG-DMR* was required for repression of *Dlk1* in both E14.5 liver and yolk sac tissues (Figure 4), a result consistent with the previous analysis of *ΔIG^MAT^* mutants (Figure 1B; 31, 44). However, while our experiments showed a loss of *Gtl2* expression in *ΔIG^MAT^* liver samples (Figure 4A, p = 0.029, wt n=3, *ΔIG^MAT^* n=5), *Gtl2* expression was not significantly altered in *ΔIG^MAT^* yolk sac tissues (Figure 4B, wt n=4, *ΔIG^MAT^* n=4). Thus, our analysis of *ΔIG^MAT^* mutants provides additional evidence for the tissue-specific roles of the maternal *IG-DMR*, revealing that this sequence is dispensable for *Gtl2* activation in the yolk sac.

**Figure 4.**
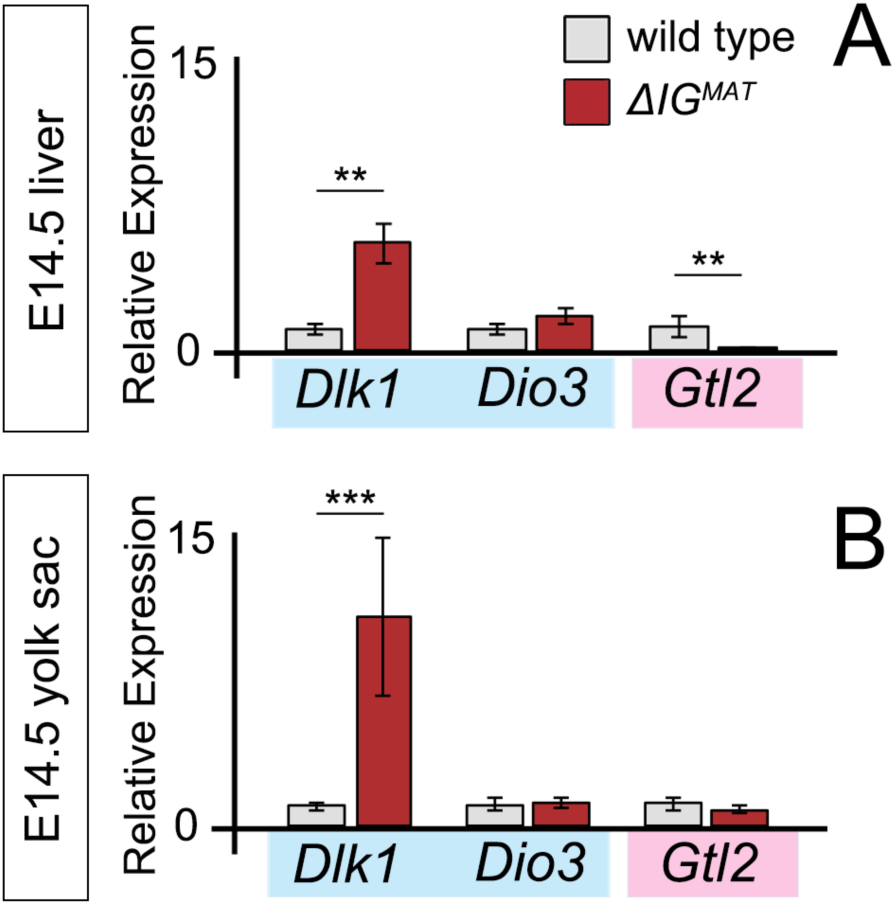
Tissue-specific effects of *ΔIG^MAT^* on *Dlk1-Dio3* gene expression. qRT-PCR of paternally-expressed (blue, *Dlk1* and *Dio3*) and maternally-expressed (pink, *Gtl2*) genes in *ΔIG^MAT^* E14.5 liver **(A)** and yolk sac **(B)** samples (red bars) as compared to wild type controls (grey bars). Expression is normalized to *β-actin* and relative to wild type littermates. For yolk sac samples, graph shows the average of four individual wild type and *ΔIG^MAT^* embryos. For liver samples, bars show the average of three wild type and five *ΔIG^MAT^* embryos. Three technical replicates were analyzed for each biological sample. Error bars represent the SEM. ** p ≤ 0.05, *** p < 0.001

### The *Dlk1-Dio3* imprinted cluster contains tissue-specific transcribed regulatory elements

Our findings that both the paternal and maternal *IG-DMR*s are required to control imprinted gene expression in a tissue-specific manner suggests that this regulatory sequence is interpreted in different ways depending on the tissue context. Notably, our analysis shows that the tissue-specific regulation of *Gtl2* expression by the *IG-DMR* does not result in tissue-specific imprinting, since *Gtl2* was expressed exclusively from the maternal allele in all tissues analyzed (Figure 3C; 18). Consequently, our results highlight that allele-specific transcriptional control is likely achieved through different mechanisms and regulatory sequences in different tissues. Specifically, our data show that deletion of the paternal *IG-DMR* has little effect on *Gtl2* expression in liver samples, but causes biallelic expression of *Gtl2* in the yolk sac, indicating that this sequence is required to repress *Gtl2* specifically in this tissue (Figure 6). Conversely, the maternal *IG-DMR* functions as an enhancer for *Gtl2* in liver tissues, but it is dispensable for *Gtl2* expression in the yolk sac (Figure 6), suggesting that additional regulatory sequences control transcription of *Gtl2* in this extraembryonic membrane.

To gain a better understanding of the regulatory landscape of the *Dlk1-Dio3* imprinted cluster, we quantified nascent transcription in tissues using RNA pol II Precision Run-On and sequencing (59) on isolated chromatin (http://biorxiv.org/lookup/doi/10.1101/185991). Because our goal was to identify tissue-specific regulatory elements that could function in *cis* to regulate allele-specific expression of *Dlk1-Dio3* imprinted genes, we performed PRO-seq on tissue samples harboring SNPs between the maternal (C57BL/6J) and paternal (CAST/EiJ) alleles at the *Dlk1-Dio3* cluster. PRO-seq data was analyzed through dREG, a machine-learning algorithm that identifies transcribed regulatory elements (TRE), including enhancers and promoters (55). Data quality analysis, including assessment of the number of sequence reads, overlap of dREG results in biological replicates, as well was the overlap of dREG peaks with enhancers and promoters (as defined by H3K27ac, H3K4me1 and H3K4me3 in ENCODE E14.5 liver datasets) is shown in Supplementary Figure 6.

PRO-seq and dREG analysis of E14.5 liver and yolk sac samples identified a total of 36 TREs within the *Dlk1-Dio3* imprinted cluster (Supplementary Figure 2). 11 TREs were common between the yolk sac and liver, 21 TREs were unique to yolk sac, and 4 TREs were unique to liver. Among the 36 identified TREs were the promoters of *Dlk1, Gtl2,* and *Dio3*, (found in both liver and yolk sac: peaks 10, 21, and 36 in Supplementary Figure 2B), the *Rtl1* promoter (found only in the liver: peak 22 in Supplementary Figure 2B), and promoters for two expressed sequence tags (ESTs, GenBank CB951180 and BU055945; Supplementary Figure 2B, peaks 23 and 29, respectively). The remaining 30 TREs showed bidirectional transcription without elongation (Supplementary Figure 3-4), a pattern indicative of open chromatin regions, including active enhancers (55). The intergenic region between *Dlk1* and *Gtl2* was particularly rich in putative regulatory regions, with 10 TREs identified by dREG (Figure 5A, Supplementary Figure 4B).

**Figure 5.**
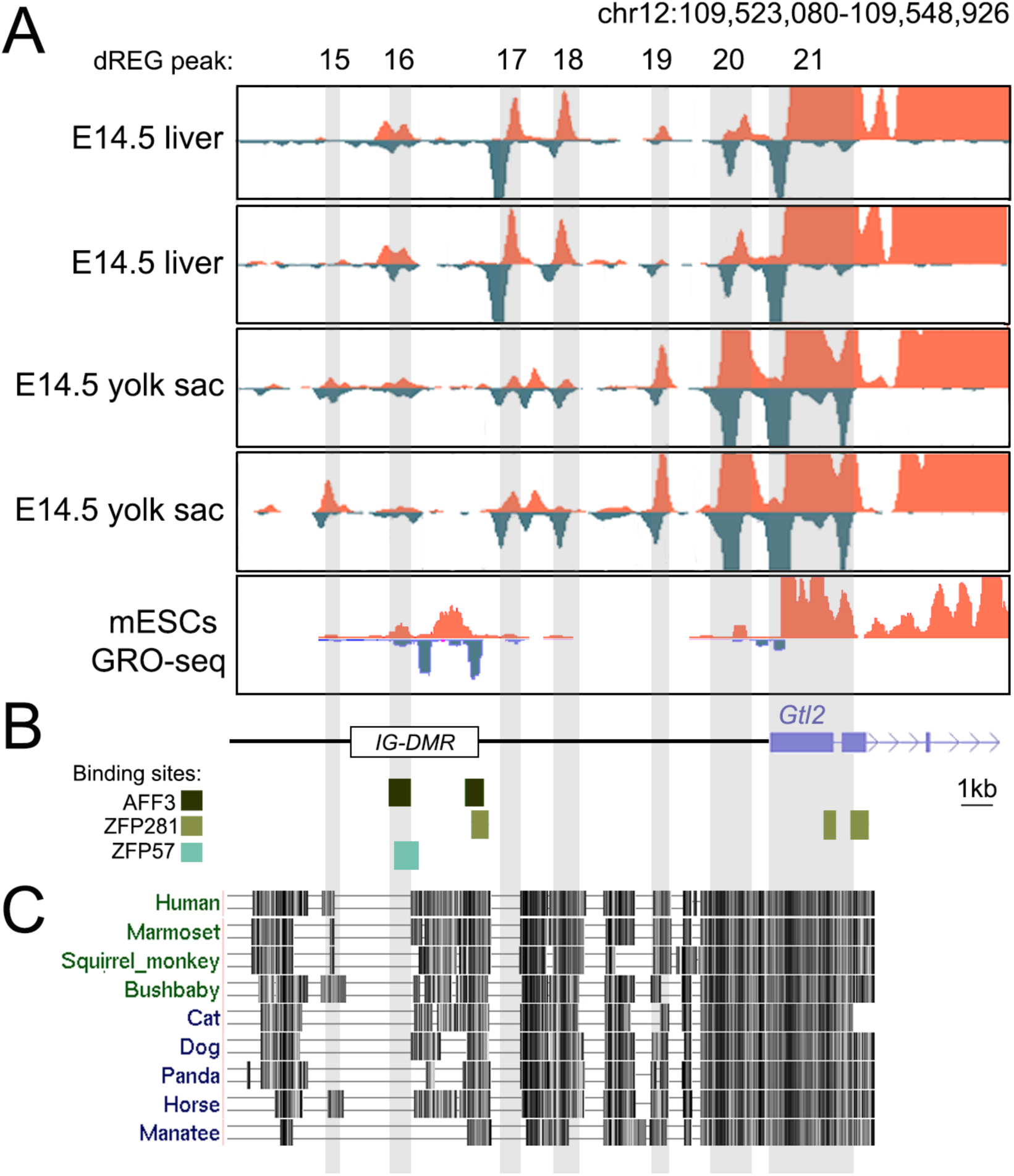
Nascent transcription at the *Dlk1-Gtl2* intergenic region. **(A)** PRO-seq in biological replicates of E14.5 liver and E14.5 yolk sac for the *Dlk1-Gtl2* intergenic region within the indicated coordinates (top, Ensembl GRCm38/UCSC mm10 coordinates). GRO-seq data from mESC (60) is shown for comparison. Blue and pink peaks highlight transcription from the negative and positive strands, respectively. The position of dREG peaks is highlighted in grey. See Supplementary Figure 2 for additional information on each dREG peak. **(B)** Map of region shown in A with the position of the 4.15 kb *IG-DMR* deletion (31), as well as AFF3, ZFP281 and ZFP57 binding sites as detected in mESCs (4, 47, 69). **(C)** UCSC genome browser conservation track between mouse and a set of placental mammals for the region shown in A.

In summary, our PRO-seq analysis identified 30 novel putative regulatory regions at the *Dlk1-Dio3* cluster, and demonstrates that the liver and yolk sac have distinct transcriptional regulatory landscapes.

### PRO-seq reveals tissue-specific transcriptional activity at the *IG-DMR*

During our analysis, we noticed that the *IG-DMR* showed a different PRO-seq profile in the liver as compared to the yolk sac (Figure 5A, peak 16). In fact, dREG analysis identified peak 16 as a liver-specific TRE, a discovery consistent with our findings that the maternal *IG-DMR* is essential for activation of *Gtl2* in the liver, but not in the yolk sac. These observations suggest that the *IG-DMR* might contain several independent regulatory sequences with tissue-specific functions. To further explore this possibility, we compared our PRO-seq findings with a published analysis of nascent transcription in mESCs using genome-wide nuclear run-on (GRO-seq; 60). This comparison revealed that mESCs have a distinct TRE profile, with nascent transcription located at different *IG-DMR* coordinates than those we observed in liver or yolk sac samples (Figure 5A). Together, these observations suggest that the *IG-DMR* contains distinct regulatory sequences that operate in different developmental contexts.

### PRO-seq identifies a maternal-specific TRE in yolk sac tissues

To determine whether the our PRO-seq and dREG datasets could identify novel TREs responsible for tissue-specific imprinting control, we searched for TREs that were expressed in an allele-specific manner by quantifying the number of PRO-seq reads containing maternal or paternal SNPs. Of the 36 TREs identified within the *Dlk1-Dio3* cluster, 25 contained annotated SNPs (Supplementary Figure 2B). The majority of these TREs (20 TREs) showed expression from both parental alleles, but 5 TREs (peaks 10, 19, 20, 21, and 22) showed monoallelic expression. We found that 3 of these 5 monoallelic TREs correspond to loci expected to have monoallelic activity, including the *Dlk1* promoter (peak 10, paternally expressed), the *Gtl2* promoter (peak 21, maternally expressed) and the *Rtl1* promoter (peak 22, paternally expressed). Interestingly, the other two monoallelic TREs mapped near the *Gtl2* promoter, in regions not previously described to contain regulatory elements (peaks 19 and 20, 5kb and 1kb from the *Gtl2* promoter, respectively). One of these TREs was expressed in both liver and yolk sac (peak 20), while the other one (peak 19, hereafter called TRE19) was yolk sac-specific (Figure 5A). While further experiments will be required to confirm whether TRE19 regulates allele-specific expression in yolk sac tissues, these results support PRO-seq as a valuable tool to identify novel regulatory regions within imprinted clusters.

### Sequence conservation suggests functional relevance of TREs for imprinting control

To assess whether the TREs identified in our PRO-seq datasets might be functionally significant, we examined the sequence conservation at the *Dlk1-Gtl2* intergenic region across several placental mammals. Interestingly, this analysis revealed that many of the TREs identified in our studies were located in areas of high sequence conservation amongst species (Figure 5C). Notably, the yolk sac-specific TRE19, TRE20, as well as the liver-specific TRE18 were all highly conserved, as was the region of the *IG-DMR* that shows mESC-specific nascent transcription. The conservation plot also revealed areas of high conservation that do not contain TREs in our dataset, nor nascent transcription in mESCs (60). These conserved regions could correspond to TREs that are only active in other tissues, or to repressive regulatory elements, which are not identified by nascent transcription. The fact that some of the TREs identified in our study are conserved across evolution suggests that these regions have critical regulatory functions and validates nascent transcription as a powerful tool to unravel the regulatory mechanisms that control imprinted gene expression.

## DISCUSSION

Our results demonstrate that imprinted gene expression at the *Dlk1-Dio3* cluster is regulated by an intricate transcriptional regulatory landscape, involving multiple regulatory sequences that are interpreted in a tissue-specific fashion. By investigating the functions of the *IG-DMR* in early embryos, liver and yolk sac, we uncovered distinct requirements for both the maternal and paternal *IG-DMRs* in each of these settings. Additionally, our analysis of nascent transcription identified 30 novel regions containing potential regulatory elements, some of which display tissue-specific activity. These results, together with our discovery that TRIM28 operates both at the *IG-DMR* and at additional regulatory sequences, challenge the view that imprinted gene expression at an individual imprinted cluster is regulated through a single mechanism across tissue types and embryonic stages. Instead, our findings reveal that multiple regulatory sequences cooperate to regulate imprinted gene expression in a tissue-specific fashion.

### After genome-wide reprogramming, TRIM28 controls imprinting through regulatory regions other than the *IG-DMR*

Our previous analysis of *Trim28* conditional mutants and hypomorphic *Trim28^chatwo^* embryos revealed that TRIM28 influences imprinted gene expression after genome-wide reprogramming through mechanisms that do not involve the maintenance of DNA methyl marks at the *IG-DMR* (50). Because TRIM28 has been described to bind to the methylated *IG-DMR* in mESCs (4), we investigated whether TRIM28 functions by binding the methylated paternal *IG-DMR* and preventing enhancer activity from this sequence. While our results show that loss of TRIM28 function causes biallelic expression of *IG-DMR RNAs*, our data indicate that this effect cannot fully explain the imprinting defects of *Trim28^chatwo^* embryos. Furthermore, epistasis experiments between *Trim28^chatwo^* and *ΔIG^PAT^* mutants indicate that TRIM28 controls imprinting after genome-wide reprogramming by mechanisms that do not involve binding to the paternal *IG-DMR*. Consequently, our results suggest that TRIM28 binds to additional regulatory regions to regulate imprinting. Published ChIP-seq experiments in ESCs have not revealed any TRIM28 binding regions at the *Dlk1-Dio3* cluster other than the *IG-DMR* (4). Nonetheless, since this published study was performed in mESCs, we cannot exclude the possibility that TRIM28 binds to other regions in different cell types. The DNA target specificity of TRIM28 has been ascribed to its ability to interact with members of the KRAB zinc finger protein family (KRAB-ZFP; 61) and a recent ChIP-exo survey of human KRAB-ZFPs in 293T cells identified several KRAB-ZFPs that bind near *Gtl2* (62). Of the 228 human KRAB-ZFPs reported in this study, 8 showed binding within 2kb of the *Gtl2* promoter (ZNF445, ZNF793, ZFP57, ZNF2, ZNF28, ZNF383, ZNF684, ZNF468), 4 bound 13kb upstream of *Gtl2* near the *IG-DMR* (ZNF257, ZNF528, ZNF445, ZNF429), and one 6kb upstream of *Gtl2* (ZNF780A) (Supplementary Figure 5). Although the results from this ChIP-exo study correspond to human KRAB-ZFPs and remain to be validated, this analysis offers several candidate locations other than the *IG-DMR* where TRIM28 could bind to regulate imprinting. While further studies will be needed to elucidate the mechanisms by which TRIM28 controls imprinting after genome-wide reprogramming, our data demonstrates that TRIM28 functions through mechanisms that do not involve *IG-DMR* binding. This finding underscores our conclusion that multiple regulatory sequences are responsible for proper imprinting control.

### Dlk1-Dio3 imprinting requires both maternal and paternal *IG-DMRs* in a tissue-specific fashion

Previous studies determined that deletion of the 4.15 kb paternal *IG-DMR* is dispensable for imprinting control (Figure 1C; 31, 44). However, our experiments in early embryos and embryonic tissues demonstrate that both the maternal and the paternal alleles of this 4.15 kb *IG-DMR* sequence are required to control imprinted gene expression. We presume that effects of the paternal 4.15 kb *IG-DMR* deletion could have been undetected in previous studies due to differences in the sensitivity of the assays used (Northern blot in ref 31; qRT-PCR in our study) or because of differences between the tissues and stages analyzed (whole E16 embryos in ref 31; individual E14.5 tissues and E8.5 embryos in our study). Interestingly, our results show that deletion of the *IG-DMR* has different effects in different tissues, as well as on specific imprinted genes. For instance, we show that deletion of the maternal *IG-DMR* has effects on *Dlk1* and *Gtl2* expression in E14.5 liver, but only disrupts *Dlk1* imprinting in E14.5 yolk sac. These tissue-specific requirements are consistent with those previously described for the maternal *IG-DMR* in embryonic versus placental tissues (44). Specifically, Lin *et al.* found that the maternal *IG-DMR* controls *Dlk1* imprinting in both E16 embryonic and placental tissues, but is only required for *Gtl2* expression in embryos and not in placentas, a tissue subject to similar imprinting control as the yolk sac (36, 63). Our results expand these observations by providing evidence that deletion of the paternal *IG-DMR* also has different effects in different tissues. Namely, we find that the paternal *IG-DMR* controls *Gtl2* expression in the yolk sac, but not in the liver (Figure 6B-C). Notably, while the 4.15 kb maternal *IG-DMR* deletion is embryonic lethal by E16, paternal *IG-DMR* deletion mutants are viable and fertile (31). The fact that imprinting defects in *Δ*IG^*PAT*^ mutants are compatible with embryonic survival is likely due to the tissue-specific effects of this regulatory sequence and highlights that some tissues can tolerate differences in the total amount of *Dlk1*, *Dio3* and *Gtl2*.

Tissue-specific imprinting effects have been previously described for some of the genes at the *Dlk1-Dio3* cluster. *Dlk1* shows biallelic expression in astrocytes (35), a relaxation of imprinting in the liver (36) and imprinting reversal in ESCs (24), while *Dio3* is biallelically expressed in several tissues (37). Additionally, the levels of expression of these genes are variable across tissues and developmental stages. For instance, *Dlk1* is expressed at high levels in many embryonic and extraembryonic tissues, including lung and liver, while expression is downregulated in most adult tissues postnatally (34). Also worth considering is the fact that all genes in the cluster are not expressed simultaneously in the same tissues and/or developmental stages (34, 40), arguing that expression of each gene in the *Dlk1-Dio3* cluster is regulated by separate regulatory sequences (64). Here, we show that expression of *Gtl2*, which is not known to be subject to tissue-imprinting effects (18, 34) and is expressed from the maternal allele in all the tissues we analyzed (Figure 3C), is disrupted by deletion of either the maternal or paternal *IG-DMRs* in a tissue-specific fashion. Therefore, our findings provide evidence that imprinted maternal *Gtl2* expression is regulated by distinct mechanisms and regulatory sequences in different tissues.

### The *IG-DMR* has tissue-specific regulatory functions

By showing that the deletion of either the maternal or paternal *IG-DMRs* has different tissue-specific effects on *Dlk1*, *Dio3* and *Gtl2* expression, our results demonstrate that the *IG-DMR* regulates imprinting through different molecular mechanisms. Tissue-specific requirements for the germline DMR have been previously described for other imprinted clusters, including the *Igf2-H19* loci (65-68). In this case, tissue-specific effects of ICR deletions have been partially attributed to tissue-specific regulatory elements that reside within the ICR, to tissue-specific modifications in the transcription factors known to bind the ICR, or to disruptions in the long range chromatin interactions that take place between tissue-specific enhancers, intergenic silencers and gene promoters (65, 68). Although these general mechanisms might also help explain the tissue-specific imprinting effects caused by deletion of the *IG-DMR*, findings at the *H19-Igf2* locus are difficult to extrapolate, given the drastic differences in imprinting control between these two imprinted clusters. For instance, the maternal unmethylated ICR at the *H19-Igf2* loci binds the chromatin insulator CCCTC-binding factor (CTCF) and functions as a boundary element that restricts the *cis* interaction of shared mesoderm and endoderm enhancers with the *Igf2* promoter (6, 8). However, CTCF binding or insulator activity have not been described to play major roles in regulating *Dlk1*-*Gtl2* imprinting (7) and expression of *Dlk1* and *Gtl2* is for the most part controlled by independent regulatory elements, rather than common enhancers (64).

While our data cannot exclude the possibility that, similar to the *H19-Igf2* cluster, some of the tissue-specific effects of the *IG-DMR* deletion might be the consequence of long-range disruptions in chromatin conformation, our results provide support for the hypothesis that the regulatory sequences located at the *IG-DMR* are interpreted in a tissue-specific fashion. In particular, we found that the PRO-seq profile of the *IG-DMR* is different between E14.5 liver, E14.5 yolk sac and a previous GRO-seq study performed in mESCs (60). Interestingly, this GRO-seq study shows mESC-specific enhancer activity at the 3’end of the *IG-DMR*, in an area where the transcriptional regulators AFF3 and ZFP281 have been described to bind the *IG-DMR* and regulate *IG-DMR* RNA expression (Figure 5B; 47, 69). Together, these findings suggest that within the *IG-DMR*, as defined by the 4.15 kb deleted in *ΔIG* mutants (31), there are several regulatory sequences with the ability to recruit a variety of transcription factors, some of which are likely expressed only in certain cell types.

### Multiple regulatory sequences are required for proper imprinting control

By showing that TRIM28 controls imprinting through regulatory sequences other than the *IG-DMR*, and that the maternal *IG-DMR* is dispensable for *Gtl2* expression in the yolk sac, our results provide evidence that multiple regulatory sequences are required for proper imprinting control at the *Dlk1-Dio3* cluster. Therefore, the identification of these regulatory sequences is critical to achieve a better understanding of the mechanisms of imprinting control at this locus. Analysis of transgenic mice bearing bacterial artificial chromosomes (BACs) has provided some clues about the location of regulatory sequences. These studies show that enhancers for expression of *Gtl2* in embryo, placenta and adult brain are localized within a BAC transgene spanning from 3.5kb upstream of *Dlk1* to 59 kb downstream of *Gtl2* (34), and that most of the regulatory sequences for *Dlk1* are located in the region contained from 41 kb upstream to 36 kb downstream of the *Dlk1* coding region (64, 70). While these studies provide evidence that tissue-specific regulation of *Dlk1* and *Gtl2* relies on separate regulatory elements, the precise location of all tissue-specific enhancers, as well as the mechanisms by which these sequences coordinate with the *IG-DMR* to ensure allele-specific expression remain to be discovered. By using PRO-seq and dREG to compare nascent transcription between E14.5 liver and yolk sac tissues, we identified 30 novel TREs at the *Dlk1*-*Gtl2* cluster. Although more experiments will be needed to confirm the regulatory roles of these regions, our finding that many TREs map to areas of high evolutionary conservation across mammalian species suggest that they have critical regulatory functions.

In conclusion, our results demonstrate that gene expression at the *Dlk1-Dio3* cluster is controlled by multiple regulatory sequences that exert imprinting control in a tissue-specific manner. Additionally, our analysis of nascent transcription highlights the power of this technique to uncover the regulatory mechanisms that control imprinted gene expression.

## DATA AVAILABILITY

PRO-seq data was deposited in NCBI’s Gene Expression Omnibus (71) and are accessible through GEO accession number GSE119882 (https://www.ncbi.nlm.nih.gov/geo/query/acc.cgi?acc=GSE119882).

## SUPPLEMENTARY DATA

This manuscript has 6 Supplementary Figures and 1 Supplementary Table.

## CONFLICT OF INTEREST

The authors declare no conflicts of interest.

## FUNDING

This work was supported by the National Science Foundation (IOS-1020878 to MJGG); and by a training grant from the National Institutes of Health (5T32HD057854 to KAA through the Research and Career Training in Vertebrate Developmental Genomics at Cornell University).

## ACKNOWLEDGEMENTS

We thank Marisa Bartolomei, Joanne Thorvaldsen, Aimee Juan, Marcos Simões-Costa, Paul Soloway, Roman Spektor and members of the laboratory for helpful discussions and comments on the manuscript; Shau-Ping Lin and Anne Ferguson-Smith for the *IG-DMR* deletion mutants; Digbijay Mahat and John Lis for technical assistance with PRO-seq experiments; Charles Danko for help with dREG analysis and Cornell’s CARE staff for outstanding mouse husbandry and care.

---

**Supplementary Figure S1.**
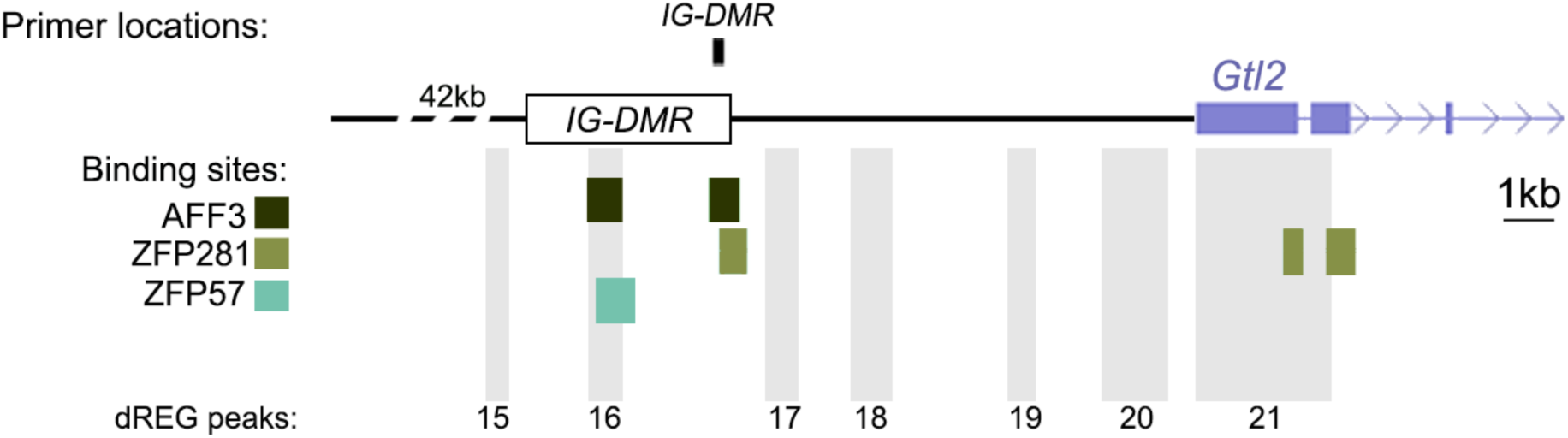
Location of regions analyzed. Region numbers match primer names in Supplementary Table 1. The positions of the *IG-DMR*, binding sites for AFF3, ZFP281 and ZFP57, as well as dREG peaks identified in our study, are indicated.

**Supplementary Figure S2.**
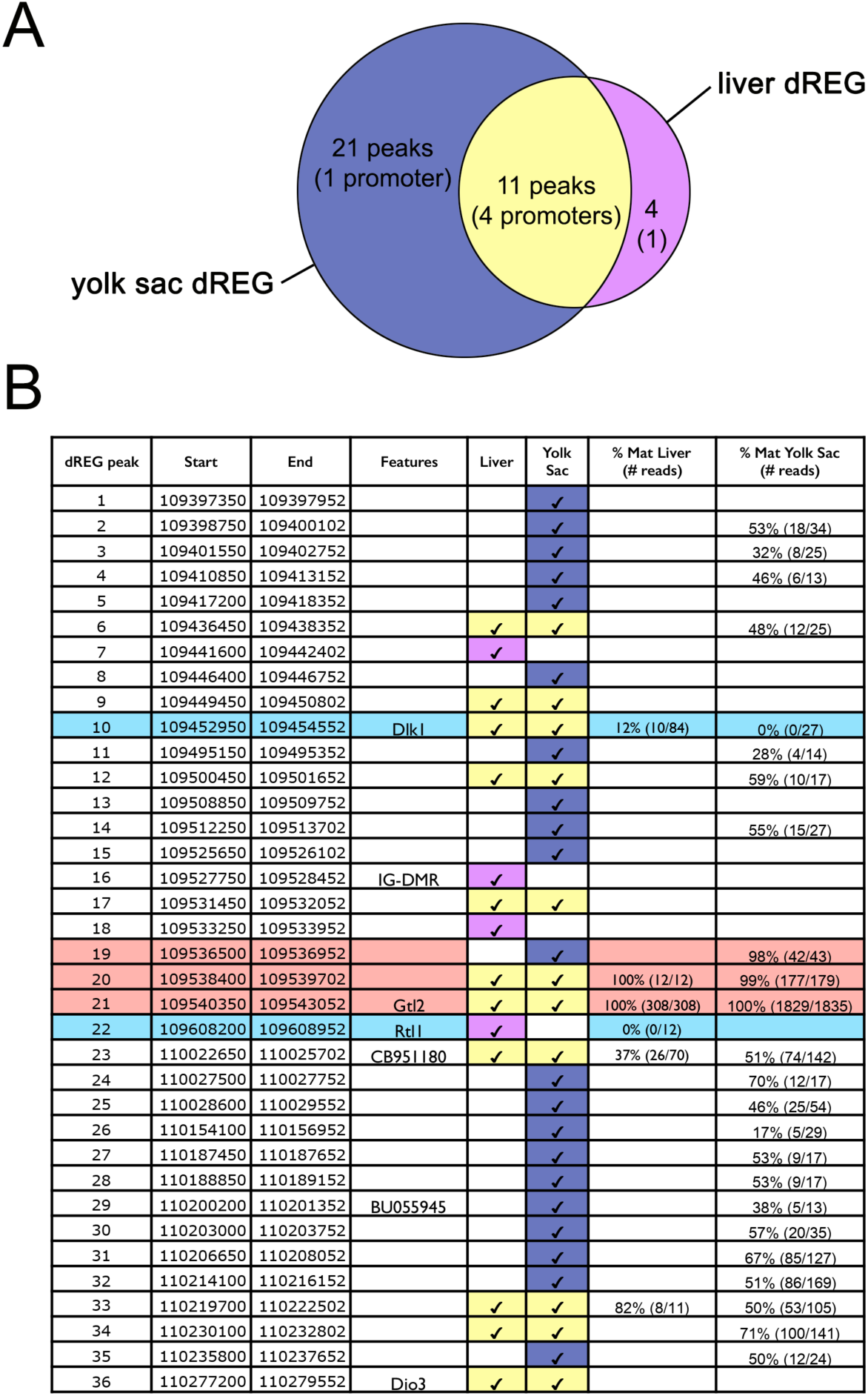
Nascent transcription dREG analysis of the *Dlk1-Dio3* imprinting cluster. **(A)** Venn diagram of overlap between dREG peaks in E14.5 liver and yolk sac tissues. **(B)** Table of all dREG peaks found within the *Dlk1-Dio3* imprinted cluster (Ensembl GRCm38/UCSC mm10 coordinates). The genomic location, known features, as well as tissue-specific and allele-specific transcriptional activities are indicated in different columns. Color highlights indicate maternal (pink), paternal (blue), liver (lilac) and yolk sac transcription (dark blue), as well as transcription in both liver and yolk sac (yellow). For dREG peaks containing SNPs, we quantified the percentage of PRO-seq reads showing maternal transcription (last two columns). Number in parentheses correspond to the total number of maternal reads over the total number of reads at that location (sum of both biological replicates).

**Supplementary Figure S3.**
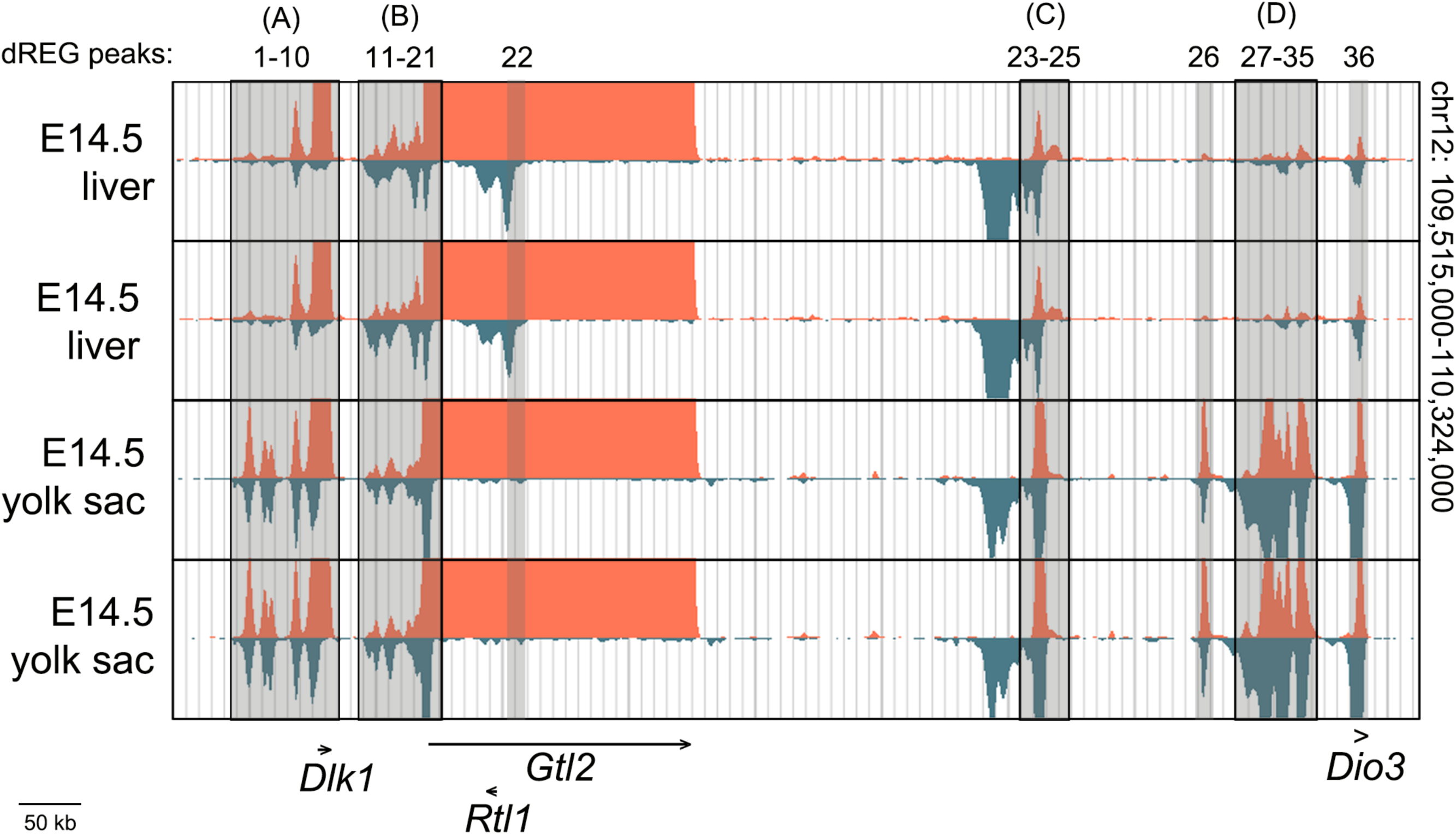
PRO-seq tracks of the *Dlk1-Dio3* cluster. Areas containing dREG peaks are highlighted in grey. Transcription from the negative and positive strands is shown in blue and red respectively. Letters between brackets indicate areas magnified in the respective panels of Supplementary Figure 4.

**Supplementary Figure S4.**
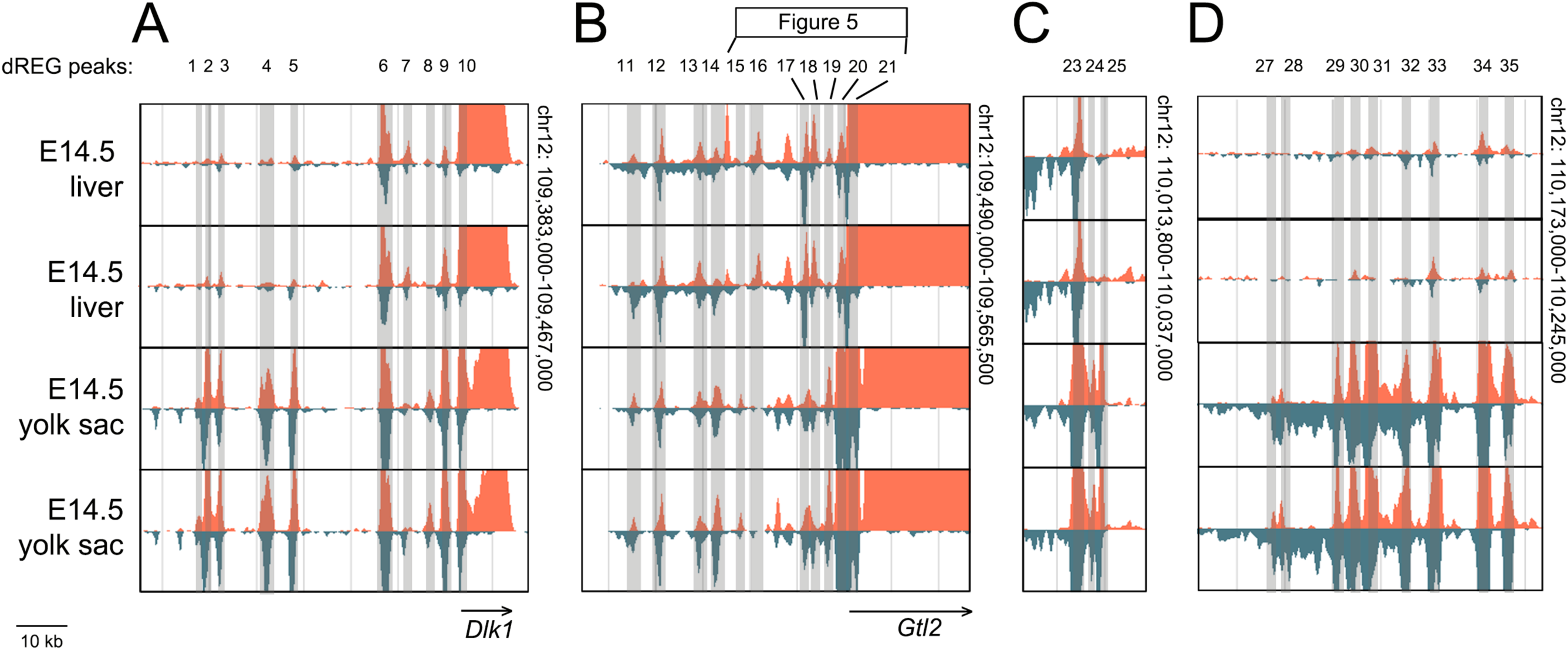
PRO-seq tracks of different areas of the *Dlk1-Dio3* cluster. Higher magnification views of areas highlighted in Supplementary Figure 3 (coordinates indicated at margin). Areas containing dREG peaks are highlighted in grey. Transcription from the negative and positive strands is shown in blue and red respectively.

**Supplementary Figure S5.**
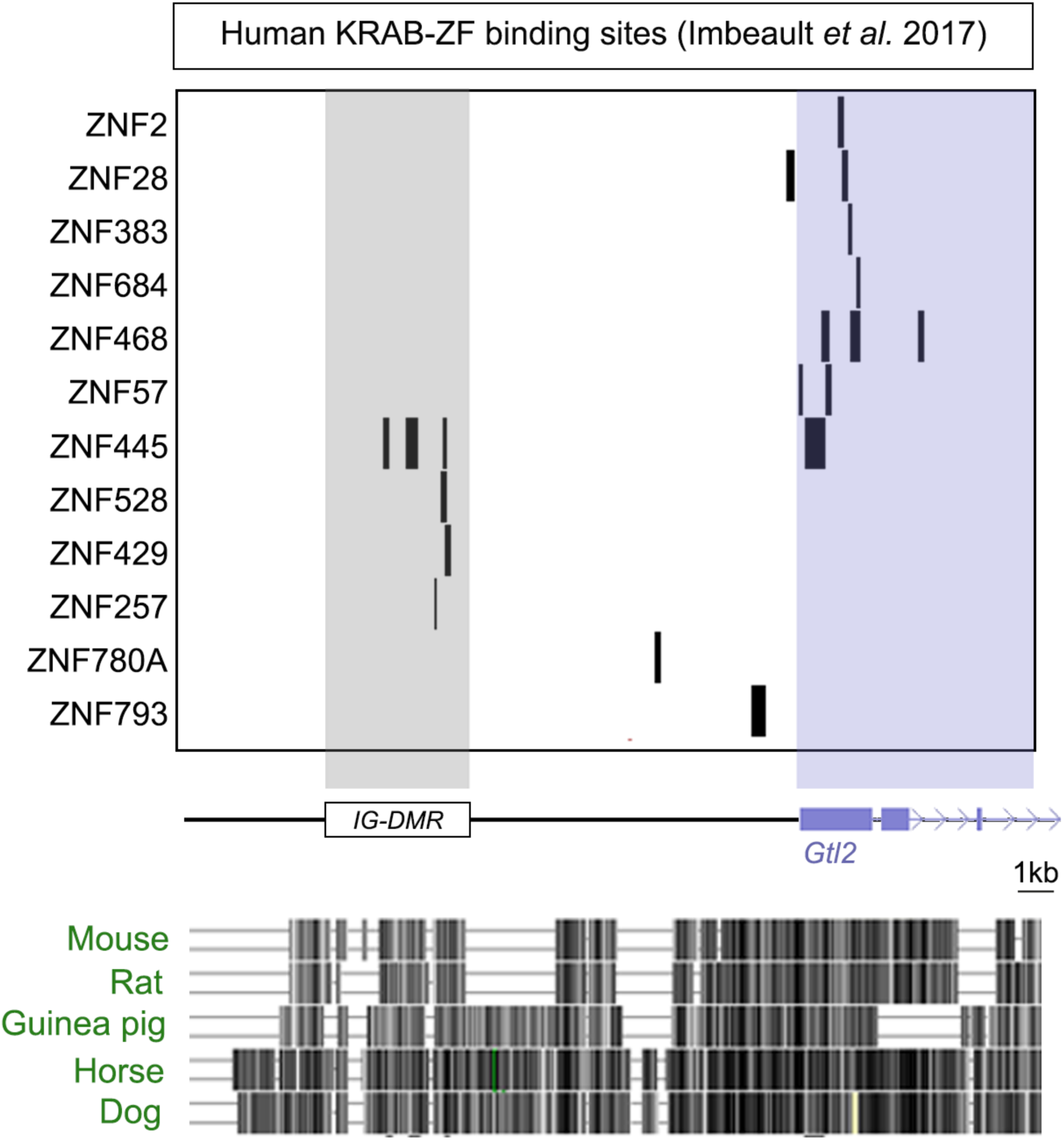
Identification of human KRAB Zinc Finger proteins at the *Dlk1-Gtl2* intergenic region. Map of the *Dlk1-Gtl2* intergenic region showing the locations bound by the indicated human KRAB domain proteins, as identified through ChIP studies in (Imbeault and Trono, 2014). UCSC genome browser conservation of the region with mouse and other placental mammals is shown below.

**Supplementary Figure S6.**
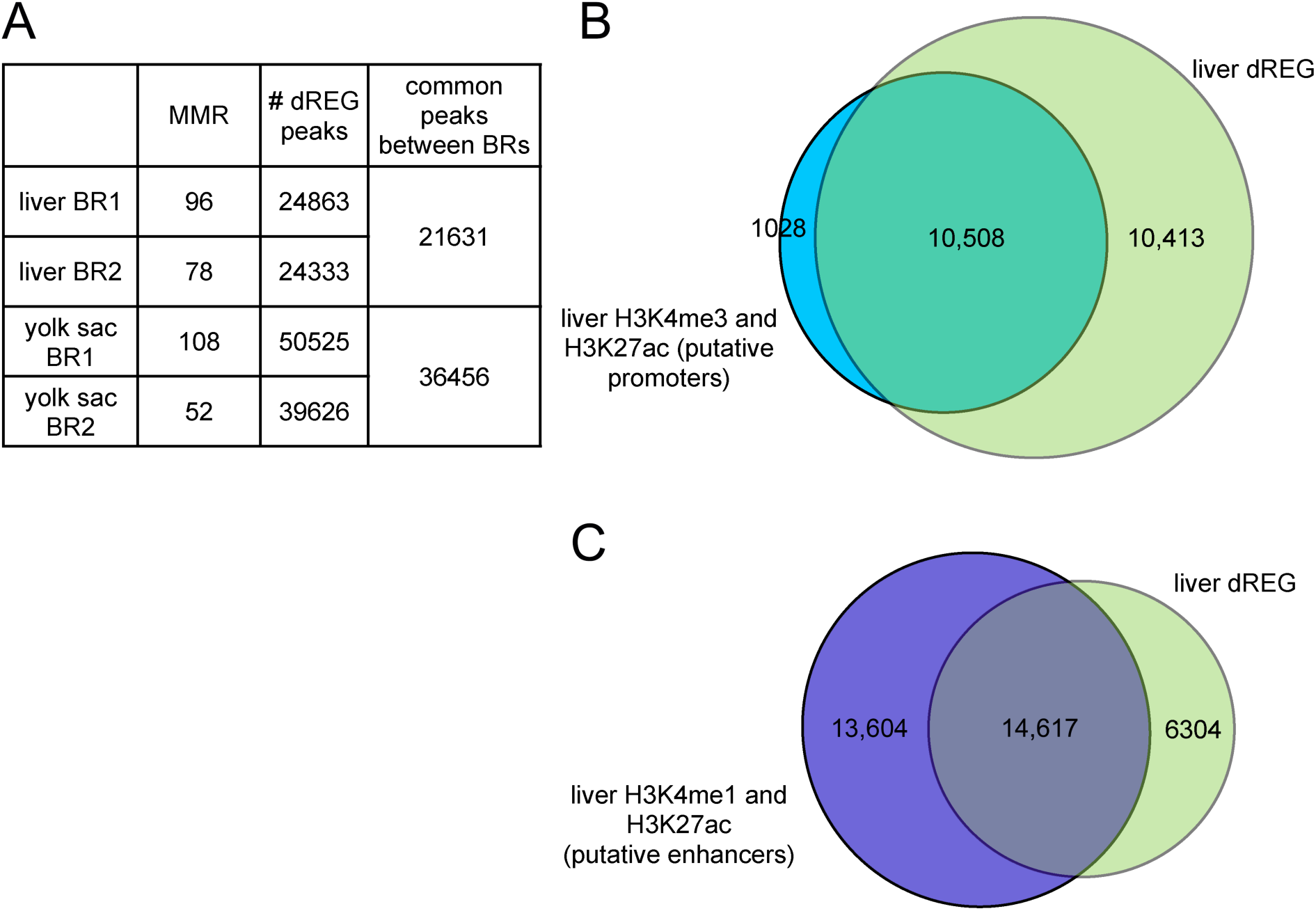
Overlap of dREG peaks with enhancers and promoters, as defined by ENCODE histone modification data. **(A)** Summary table of the number of million mapped reads (MMR), number of dREG peaks in two biological replicates (BR1 and BR2), and number of dREG peaks that are common between biological replicates. Genome-wide comparison of liver dREG analysis to ENCODE histone ChIP data showed strong overlap between identified TREs and putative promoters in E14.5 liver (91% of peaks with H3K27ac and H3K4me3 overlap with liver TREs) **(B)**, as well as substantial overlap between identified TREs and putative enhancers (51% of peaks with H3K27ac and H3K4me1 overlap with liver TREs) **(C)**. Overlap between dREG and ENCODE data was calculated using the bedtools intersect tool. dREG peaks were converted to mm9 using the liftover tool (UCSC genome browser) in order to overlap with mm9 ENCODE data (only 20,921 liver dREG peaks were successfully converted).

**Supplementary Table 1.**
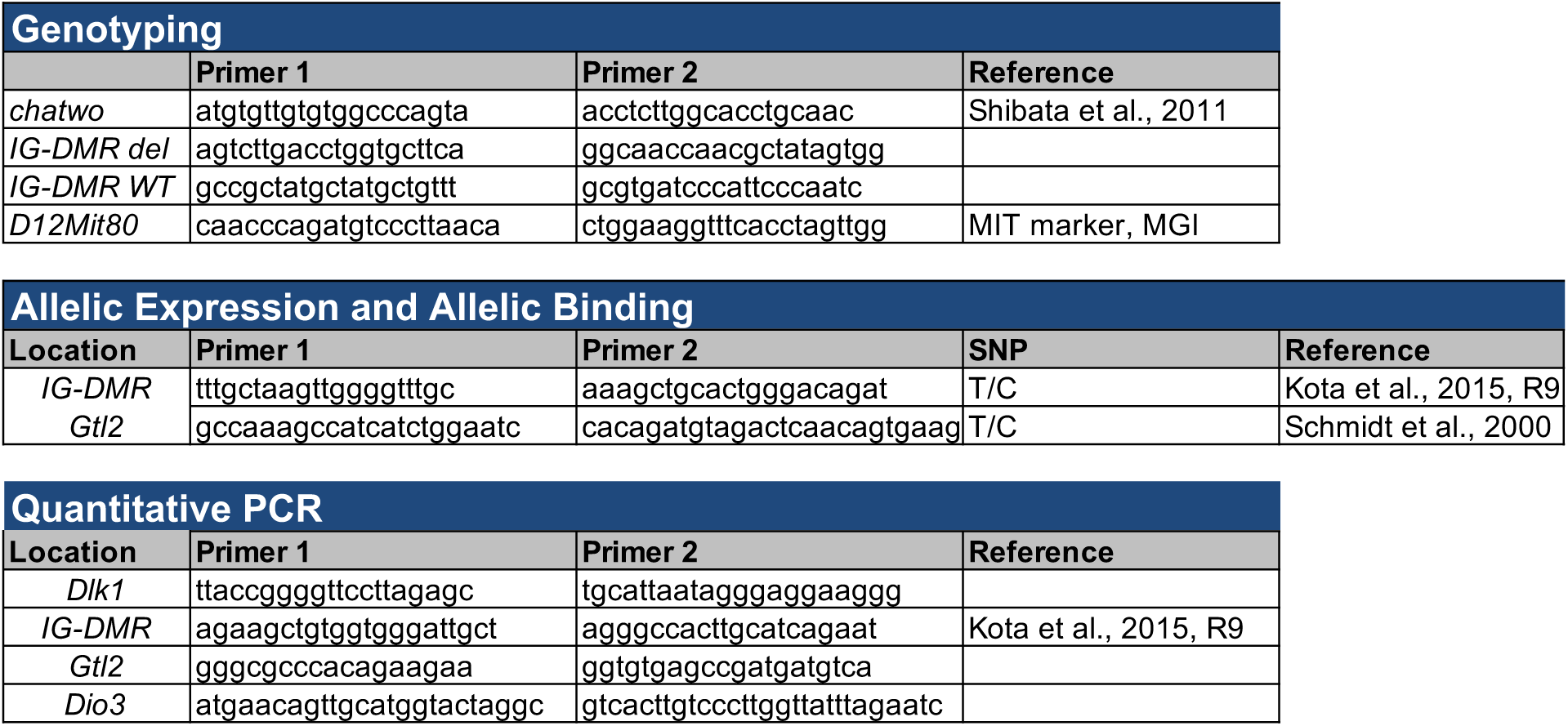
Sequences of primers used.

## References

1. Lee,J.T. and Bartolomei,M.S. (2013) X-inactivation, imprinting, and long noncoding RNAs in health and disease. Cell, 152, 1308–1323.

2. Stadtfeld,M., Apostolou,E., Akutsu,H., Fukuda,A., Follett,P., Natesan,S., Kono,T., Shioda,T. and Hochedlinger,K. (2010) Aberrant silencing of imprinted genes on chromosome 12qF1 in mouse induced pluripotent stem cells. Nature, 465, 175–181.

3. Ferguson-Smith,A.C. (2011) Timeline: Genomic imprinting: the emergence of an epigenetic paradigm. Nature Publishing Group, 12, 565–575.

4. Quenneville,S., Verde,G., Corsinotti,A., Kapopoulou,A., Jakobsson,J., Offner,S., Baglivo,I., Pedone,P.V., Grimaldi,G., Riccio,A., et al. (2011) In Embryonic Stem Cells, ZFP57/KAP1 Recognize a Methylated Hexanucleotide to Affect Chromatin and DNA Methylation of Imprinting Control Regions. Mol. Cell, 44, 361–372.

5. Balmer,D., Arredondo,J., Samaco,R.C. and LaSalle,J.M. (2002) MECP2 mutations in Rett syndrome adversely affect lymphocyte growth, but do not affect imprinted gene expression in blood or brain. Hum. Genet., 110, 545–552.

6. Bell,A.C. and Felsenfeld,G. (2000) Methylation of a CTCF-dependent boundary controls imprinted expression of the Igf2 gene. Nature, 405, 482–485.

7. Carr,M.S., Yevtodiyenko,A., Schmidt,C.L. and Schmidt,J.V. (2007) Allele-specific histone modifications regulate expression of the Dlk1–Gtl2 imprinted domain. Genomics, 89, 280–290.

8. Hark,A.T., Schoenherr,C.J., Katz,D.J., Ingram,R.S., Levorse,J.M. and Tilghman,S.M. (2000) CTCF mediates methylation-sensitive enhancer-blocking activity at the H19/Igf2 locus. Nature, 405, 486–489.

9. Kanduri,C., Pant,V., Loukinov,D., Pugacheva,E., Qi,C.F., Wolffe,A., Ohlsson,R. and Lobanenkov,V.V. (2000) Functional association of CTCF with the insulator upstream of the H19 gene is parent of origin-specific and methylation-sensitive. Curr. Biol., 10, 853–856.

10. Samaco,R.C., Hogart,A. and LaSalle,J.M. (2005) Epigenetic overlap in autism-spectrum neurodevelopmental disorders: MECP2 deficiency causes reduced expression of UBE3A and GABRB3. Human Molecular Genetics, 14, 483–492.

11. Szabó,P., Tang,S.H., Rentsendorj,A., Pfeifer,G.P. and Mann,J.R. (2000) Maternal-specific footprints at putative CTCF sites in the H19 imprinting control region give evidence for insulator function. Curr. Biol., 10, 607–610.

12. Szabo,P.E., Tang,S.H.E., Silva,F.J., Tsark,W.M.K. and Mann,J.R. (2004) Role of CTCF Binding Sites in the Igf2/H19 Imprinting Control Region. Molecular and Cellular Biology, 24, 4791–4800.

13. Filion,G.J.P., Zhenilo,S., Salozhin,S., Yamada,D., Prokhortchouk,E. and Defossez,P.-A. (2006) A family of human zinc finger proteins that bind methylated DNA and repress transcription. Molecular and Cellular Biology, 26, 169–181.

14. Hendrich,B., Guy,J., Ramsahoye,B., Wilson,V.A. and Bird,A. (2001) Closely related proteins MBD2 and MBD3 play distinctive but interacting roles in mouse development. Genes Dev., 15, 710–723.

15. Monnier,P., Martinet,C., Pontis,J., Stancheva,I., Ait-Si-Ali,S. and Dandolo,L. (2013) H19 lncRNA controls gene expression of the Imprinted Gene Network by recruiting MBD1. Proceedings of the National Academy of Sciences, 110, 20693–20698.

16. Prickett,A.R. and Oakey,R.J. (2012) A survey of tissue-specific genomic imprinting in mammals. Molecular Genetics and Genomics, 287, 621–630.

17. Hudson,Q.J., Kulinski,T.M., Huetter,S.P. and Barlow,D.P. (2010) Genomic imprinting mechanisms in embryonic and extraembryonic mouse tissues. Heredity, 105, 45–56.

18. Baran,Y., Subramaniam,M., Biton,A., Tukiainen,T., Tsang,E.K., Rivas,M.A., Pirinen,M., Gutierrez-Arcelus,M., Smith,K.S., Kukurba,K.R., et al. (2015) The landscape of genomic imprinting across diverse adult human tissues. Genome Research, 25, 927–936.

19. Szabo,P.E. and Mann,J.R. (1995) Allele-specific expression and total expression levels of imprinted genes during early mouse development: implications for imprinting mechanisms. Genes Dev., 9, 3097–3108.

20. Lerchner,W. and Barlow,D.P. (1997) Paternal repression of the imprinted mouse Igf2r locus occurs during implantation and is stable in all tissues of the post-implantation mouse embryo. Mech. Dev., 61, 141–149.

21. Yamasaki,Y., Kayashima,T., Soejima,H., Kinoshita,A., Yoshiura,K.-I., Matsumoto,N., Ohta,T., Urano,T., Masuzaki,H., Ishimaru,T., et al. (2005) Neuron-specific relaxation of Igf2r imprinting is associated with neuron-specific histone modifications and lack of its antisense transcript Air. Human Molecular Genetics, 14, 2511–2520.

22. Arnaud,P., Monk,D., Hitchins,M., Gordon,E., Dean,W., Beechey,C.V., Peters,J., Craigen,W., Preece,M., Stanier,P., et al. (2003) Conserved methylation imprints in the human and mouse GRB10 genes with divergent allelic expression suggests differential reading of the same mark. Human Molecular Genetics, 12, 1005–1019.

23. Sanz,L.A., Chamberlain,S., Sabourin,J.-C., Henckel,A., Magnuson,T., Hugnot,J.-P., Feil,R. and Arnaud,P. (2008) A mono-allelic bivalent chromatin domain controls tissue-specific imprinting at Grb10. EMBO J., 27, 2523–2532.

24. Kota,S.K., Llères,D., Bouschet,T., Hirasawa,R., Marchand,A., Begon-Pescia,C., Sanli,I., Arnaud,P., Journot,L., Girardot,M., et al. (2014) ICR Noncoding RNA Expression Controls Imprinting and DNA Replication at the Dlk1-Dio3 Domain. Developmental Cell, 31, 19–33.

25. Hu,J.F., Oruganti,H., Vu,T.H. and Hoffman,A.R. (1998) Tissue-specific imprinting of the mouse insulin-like growth factor II receptor gene correlates with differential allele-specific DNA methylation. Mol. Endocrinol., 12, 220–232.

26. Williamson,C.M., Ball,S.T., Nottingham,W.T., Skinner,J.A., Plagge,A., Turner,M.D., Powles,N., Hough,T., Papworth,D., Fraser,W.D., et al. (2004) A cis-acting control region is required exclusively for the tissue-specific imprinting of Gnas. Nat Genet, 36, 894–899.

27. Yamasaki-Ishizaki,Y., Kayashima,T., Mapendano,C.K., Soejima,H., Ohta,T., Masuzaki,H., Kinoshita,A., Urano,T., Yoshiura,K.-I., Matsumoto,N., et al. (2007) Role of DNA methylation and histone H3 lysine 27 methylation in tissue-specific imprinting of mouse Grb10. Molecular and Cellular Biology, 27, 732–742.

28. Valleley,E.M., Cordery,S.F. and Bonthron,D.T. (2007) Tissue-specific imprinting of the ZAC/PLAGL1 tumour suppressor gene results from variable utilization of monoallelic and biallelic promoters. Human Molecular Genetics, 16, 972–981.

29. Weinstein,L.S., Liu,J., Sakamoto,A., Xie,T. and Chen,M. (2004) Minireview: GNAS: normal and abnormal functions. Endocrinology, 145, 5459–5464.

30. John,R.M. and Lefebvre,L. (2011) Developmental regulation of somatic imprints. Differentiation, 81, 270–280.

31. Lin,S.-P., Youngson,N., Takada,S., Seitz,H., Reik,W., Paulsen,M., Cavaille,J. and Ferguson-Smith,A.C. (2003) Asymmetric regulation of imprinting on the maternal and paternal chromosomes at the Dlk1-Gtl2 imprinted cluster on mouse chromosome 12. Nat wGenet, 35, 97–102.

32. Steshina,E.Y., Carr,M.S., Glick,E.A., Yevtodiyenko,A., Appelbe,O.K. and Schmidt,J.V. (2006) Loss of imprinting at the Dlk1-Gtl2 locus caused by insertional mutagenesis in the Gtl2 5’ region. BMC Genet., 7, 44.

33. da Rocha,S.T., Edwards,C.A., Ito,M., Ogata,T. and Ferguson-Smith,A.C. (2008) Genomic imprinting at the mammalian Dlk1-Dio3 domain. Trends in Genetics, 24, 306–316.

34. Yevtodiyenko,A., Steshina,E.Y., Farner,S.C., Levorse,J.M. and Schmidt,J.V. (2004) A 178- kb BAC transgene imprints the mouse Gtl2 gene and localizes tissue-specific regulatory elements. Genomics, 84, 277–287.

35. Ferrón,S.R., Charalambous,M., Radford,E., McEwen,K., Wildner,H., Hind,E., Morante-Redolat,J.M., Laborda,J., Guillemot,F., Bauer,S.R., et al. (2011) Postnatal loss of Dlk1 imprinting in stem cells and niche astrocytes regulates neurogenesis. Nature, 475, 381–385.

36. Swanzey,E. and Stadtfeld,M. (2016) A reporter model to visualize imprinting stability at the Dlk1 locus during mouse development and in pluripotent cells. Development, 10.1242/dev.138255.supplemental.

37. Martinez,M.E., Charalambous,M., Saferali,A., Fiering,S., Naumova,A.K., St Germain,D., Ferguson-Smith,A.C. and Hernandez,A. (2014) Genomic imprinting variations in the mouse type 3 deiodinase gene between tissues and brain regions. Mol. Endocrinol., 28, 1875–1886.

38. Schmidt,J.V., Matteson,P.G., Jones,B.K., Guan,X.J. and Tilghman,S.M. (2000) The Dlk1 and Gtl2 genes are linked and reciprocally imprinted. Genes Dev., 14, 1997–2002.

39. Yevtodiyenko,A. and Schmidt,J.V. (2006) Dlk1 expression marks developing endothelium and sites of branching morphogenesis in the mouse embryo and placenta. Dev. Dyn., 235, 1115–1123.

40. Schuster-Gossler,K., Bilinski,P., Sado,T., Ferguson-Smith,A. and Gossler,A. (1998) The mouse Gtl2 gene is differentially expressed during embryonic development, encodes multiple alternatively spliced transcripts, and may act as an RNA. Dev. Dyn., 212, 214–228.

41. Seibt,J., Armant,O., Le Digarcher,A., Castro,D., Ramesh,V., Journot,L., Guillemot,F., Vanderhaeghen,P. and Bouschet,T. (2012) Expression at the Imprinted Dlk1-Gtl2 Locus Is Regulated by Proneural Genes in the Developing Telencephalon. PLoS ONE, 7, e48675.

42. Floridon,C., Jensen,C.H., Thorsen,P., Nielsen,O., Sunde,L., Westergaard,J.G., Thomsen,S.G. and Teisner,B. (2000) Does fetal antigen 1 (FA1) identify cells with regenerative, endocrine and neuroendocrine potentials? A study of FA1 in embryonic, fetal, and placental tissue and in maternal circulation. Differentiation, 66, 49–59.

43. Takada,S., Tevendale,M., Baker,J., Georgiades,P., Campbell,E., Freeman,T., Johnson,M.H., Paulsen,M. and Ferguson-Smith,A.C. (2000) Delta-like and gtl2 are reciprocally expressed, differentially methylated linked imprinted genes on mouse chromosome 12. Curr. Biol., 10, 1135–1138.

44. Lin,S.P., Coan,P., da Rocha,S.T., Seitz,H., Cavaille,J., Teng,P.W., Takada,S. and Ferguson-Smith,A.C. (2007) Differential regulation of imprinting in the murine embryo and placenta by the Dlk1-Dio3 imprinting control region. Development, 134, 417–426.

45. Li,X., Ito,M., Zhou,F., Youngson,N., Zuo,X., Leder,P. and Ferguson-Smith,A.C. (2008) A Maternal-Zygotic Effect Gene, Zfp57, Maintains Both Maternal and Paternal Imprints. Developmental Cell, 15, 547–557.

46. Das,P.P., Hendrix,D.A., Apostolou,E., Buchner,A.H., Canver,M.C., Beyaz,S., Ljuboja,D., Kuintzle,R., Kim,W., Karnik,R., et al. (2015) PRC2 Is Required to Maintain Expression of the Maternal Gtl2-Rian-Mirg Locus by Preventing De Novo DNA Methylation in Mouse Embryonic Stem Cells. CellReports, 12, 1456–1470.

47. Luo,Z., Lin,C., Woodfin,A.R., Bartom,E.T., Gao,X., Smith,E.R. and Shilatifard,A. (2016) Regulation of the imprinted Dlk1-Dio3locus by allele-specific enhancer activity. Genes Dev., 30, 92–101.

48. Saito,T., Hara,S., Kato,T., Tamano,M., Muramatsu,A., Asahara,H. and Takada,S. (2018) A tandem repeat array in IG-DMR is essential for imprinting of paternal allele at the Dlk1-Dio3 domain during embryonic development. Human Molecular Genetics, 10.1093/hmg/ddy235.

49. Zuo,X., Sheng,J., Lau,H.T., McDonald,C.M., Andrade,M., Cullen,D.E., Bell,F.T., Iacovino,M., Kyba,M., Xu,G., et al. (2012) Zinc Finger Protein ZFP57 Requires Its Co-factor to Recruit DNA Methyltransferases and Maintains DNA Methylation Imprint in Embryonic Stem Cells via Its Transcriptional Repression Domain. Journal of Biological Chemistry, 287, 2107–2118.

50. Alexander,K.A., Wang,X., Shibata,M., Clark,A.G. and Garcia-Garcia,M.J. (2015) TRIM28 Controls Genomic Imprinting through Distinct Mechanisms during and after Early Genome-wide Reprogramming. CellReports, 13, 1194–1205.

51. Messerschmidt,D.M., Knowles,B.B. and Solter,D. (2014) DNA methylation dynamics during epigenetic reprogramming in the germline and preimplantation embryos. Genes Dev., 28, 812–828.

52. Shibata,M., Blauvelt,K.E., Liem,K.F. and Garcia-Garcia,M.J. (2011) TRIM28 is required by the mouse KRAB domain protein ZFP568 to control convergent extension and morphogenesis of extra-embryonic tissues. Development, 138, 5333–5343.

53. Ge,B., Gurd,S., Gaudin,T., Dore,C., Lepage,P., Harmsen,E., Hudson,T.J. and Pastinen,T. (2005) Survey of allelic expression using EST mining. Genome Research, 15, 1584–1591.

54. Mahat,D.B., Kwak,H., Booth,G.T., Jonkers,I.H., Danko,C.G., Patel,R.K., Waters,C.T., Munson,K., Core,L.J. and Lis,J.T. (2016) Base-pair-resolution genome-wide mapping of active RNA polymerases using precision nuclear run-on (PRO-seq). Nat Protoc, 11, 1455–1476.

55. Danko,C.G., Hyland,S.L., Core,L.J., Martins,A.L., Waters,C.T., Lee,H.W., Cheung,V.G., Kraus,W.L., Lis,J.T. and Siepel,A. (2015) Identification of active transcriptional regulatory elements from GRO-seq data. Nat. Methods, 12, 433–438.

56. Andersson,R., Gebhard,C., Miguel-Escalada,I., Hoof,I., Bornholdt,J., Boyd,M., Chen,Y., Zhao,X., Schmidl,C., Suzuki,T., et al. (2014) An atlas of active enhancers across human cell types and tissues. Nature, 507, 455–461.

57. Hah,N., Murakami,S., Nagari,A., Danko,C.G. and Kraus,W.L. (2013) Enhancer transcripts mark active estrogen receptor binding sites. Genome Research, 23, 1210–1223.

58. Melgar,M.F., Collins,F.S. and Sethupathy,P. (2011) Discovery of active enhancers through bidirectional expression of short transcripts. Genome Biol, 12, R113.

59. Kwak,H., Fuda,N.J., Core,L.J. and Lis,J.T. (2013) Precise maps of RNA polymerase reveal how promoters direct initiation and pausing. Science, 339, 950–953.

60. Jonkers,I., Kwak,H. and Lis,J.T. (2014) Genome-wide dynamics of Pol II elongation and its interplay with promoter proximal pausing, chromatin, and exons. Elife, 3, e02407.

61. Urrutia,R. (2003) KRAB-containing zinc-finger repressor proteins. Genome Biol, 4, 231.

62. Imbeault,M., Helleboid,P.-Y. and Trono,D. (2017) KRAB zinc-finger proteins contribute to the evolution of gene regulatory networks. Nature, 10.1038/nature21683.

63. Hudson,Q.J., Seidl,C.I.M., Kulinski,T.M., Huang,R., Warczok,K.E., Bittner,R., Bartolomei,M.S. and Barlow,D.P. (2011) Extra-embryonic-specific imprinted expression is restricted to defined lineages in the post-implantation embryo. Dev. Biol., 353, 420–431.

64. Rogers,E.D., Ramalie,J.R., McMurray,E.N. and Schmidt,J.V. (2012) Localizing Transcriptional Regulatory Elements at the Mouse Dlk1 Locus. PLoS ONE, 7, e36483.

65. Thorvaldsen,J.L., Mann,M.R.W., Nwoko,O., Duran,K.L. and Bartolomei,M.S. (2002) Analysis of sequence upstream of the endogenous H19 gene reveals elements both essential and dispensable for imprinting. Molecular and Cellular Biology, 22, 2450–2462.

66. Drewell,R.A., Brenton,J.D., Ainscough,J.F., Barton,S.C., Hilton,K.J., Arney,K.L., Dandolo,L. and Surani,M.A. (2000) Deletion of a silencer element disrupts H19 imprinting independently of a DNA methylation epigenetic switch. Development, 127, 3419–3428.

67. Kaffer,C.R., Srivastava,M., Park,K.Y., Ives,E., Hsieh,S., Batlle,J., Grinberg,A., Huang,S.P. and Pfeifer,K. (2000) A transcriptional insulator at the imprinted H19/Igf2 locus. Genes Dev., 14, 1908–1919.

68. Ideraabdullah,F.Y., Thorvaldsen,J.L., Myers,J.A. and Bartolomei,M.S. (2014) Tissue-specific insulator function at H19/Igf2 revealed by deletions at the imprinting control region. Human Molecular Genetics, 23, 6246–6259.

69. Wang,Y., Shen,Y., Dai,Q., Yang,Q., Zhang,Y., Wang,X., Xie,W., Luo,Z. and Lin,C. (2016) A permissive chromatin state regulated by ZFP281-AFF3 in controlling the imprinted Meg3 polycistron. 45, 1177–1185.

70. da Rocha,S.T., Charalambous,M., Lin,S.-P., Gutteridge,I., Ito,Y., Gray,D., Dean,W. and Ferguson-Smith,A.C. (2009) Gene dosage effects of the imprinted delta-like homologue 1 (dlk1/pref1) in development: implications for the evolution of imprinting. PLoS Genet, 5, e1000392.

71. Edgar,R., Domrachev,M. and Lash,A.E. (2002) Gene Expression Omnibus: NCBI gene expression and hybridization array data repository. Nucleic Acids Res., 30, 207–210.

72. Imbeault,M. and Trono,D. (2014) As Time Goes by: KRABs Evolveto KAP Endogenous Retroelements. Developmental Cell, 31, 257–258.

